# The metabolic enzyme EXT1 is sufficient to induce the epithelial-mesenchymal transition program in cancers

**DOI:** 10.1101/2023.01.05.522866

**Authors:** Balakrishnan Solaimuthu, Anees Khatib, Arata Hayashi, Mayur Tanna, Michal Lichtenstein, Abdelrahman Karmi, Yoav D. Shaul

## Abstract

Carcinomas often exhibit aggressive characteristics, such as enhanced migration abilities, through the execution of the epithelial-mesenchymal transition (EMT) program. Heparan sulfate (HS) is a polysaccharide expressed on the surface of aggressive cancer cells, which acts as a co-receptor to stimulate EMT-associated signaling pathways. However, despite HS’ role in cancer aggressiveness, the mechanisms governing its EMT-dependent biosynthesis remains poorly understood. Here, we characterized the HS chain elongation enzyme, exostosin glycosyltransferase 1 (EXT1), as an essential component of the EMT program. We identified an EMT-dependent expression of EXT1 and its selective upregulation in aggressive tumor subtypes and cell lines. Overexpression of EXT1 in epithelial cells is sufficient to induce HS biosynthesis, cell migration, and invasion, form tumors in mice, and activate the STAT3 pathway. Moreover, its knockout in aggressive cells significantly inhibited their EMT-associated characteristics. These findings demonstrate a cellular mechanism by which metabolic processes regulate signaling pathways to govern cell state.

## Introduction

During the past few decades, substantial progress in breast cancer diagnosis and treatment improved patient outcomes. Nevertheless, tumor metastasis and drug resistance are still significant challenges for cancer therapy ^1^. One of the potential mechanisms by which carcinomas gain these aggressive traits is through the induction of the epithelial-mesenchymal transition (EMT) program. Upon activation, this program transdifferentiates the carcinoma cells into a partially mesenchymal state^2^, leading to significant changes in cellular physiology. This program includes notable morphological changes, loss of cell-cell interactions ^3^, acquisition of migratory and invasive capabilities ^4^, and remodeling the proteoglycan landscape ^5^. Therefore, a deeper understanding of cellular mechanisms mediating this program may lead to an improved conceptualization of EMT biology and therapeutic strategies to block its progression.

The extracellular matrix (ECM) is enriched in proteoglycan (PG) and glycosaminoglycan (GSG) that form a complex network between the cells ^6^. Heparan sulfate proteoglycans (HSPGs) are the most abundant PG ^7^, ubiquitously expressed on the cell surface of diverse cell types, including tumors ^8,9^. HSPGs are composed of repeating units of glucosamine and glucuronic acid (GlcA) ^8^, which are heavily modified with various sulfate groups ^10^, resulting in highly dynamic and negatively charged structures ^9^. This modification provides binding sites for positively charged amino acids present in a broad spectrum of soluble growth factors, cytokines, membrane proteins, and components of the ECM ^11,12^. HSPG’s interaction with different ligands facilitates their binding to the receptors, which subsequently induces the activation of central signaling cascades. For example, this proteoglycan governs Janus kinase (JAK)-signal transducer and activator of transcription 3 (STAT3) pathway ^7^, which regulates cell migration ^13^, a hallmark of a partial mesenchymal state. However, the cellular mechanisms by which the EMT program regulates HSPG biosynthesis remain largely unknown.

The synthesis of HSPGs is a complex and dynamic process that involves a series of sequential reactions ^14^, including chain (synthesis) initiation, elongation, and modification ^15^. This biosynthesis machinery is localized in the Golgi apparatus ^16^ and is mediated by the five members of the exostosin family, which are type II transmembrane proteins. This family includes; exostosin glycosyltransferase 1 (EXT1), EXT2, and three EXT-like EXTL1, EXTL2, and EXTL3 ^17^. The EXT-like enzymes catalyze the initial step of the chain elongation ^9^, which involves the addition of β1,4-linked N-acetylgalactosamine (GalNAc) to the forming HS backbone ^17^. Subsequently, EXT1 and EXT2 form a stable complex, which adds a diverse number of GlcNAc and GlcA to the initial chain ^18,19^. The importance of these enzymes to HS synthesis is evidenced by EXT1 or EXT2 mutations, which result in hereditary multiple exostoses (HME) ^20^. HME is a rare skeletal disorder characterized by multiple bony protuberances called exostoses that could transform into malignant chondrosarcoma ^21,22^. Here we found that EXT1, through HSPG biosynthesis, is an essential component of the EMT program. Specifically, we demonstrated a direct correlation between EXT1 expression and EMT-dependent cancer aggressiveness phenotypes. Therefore, we revealed the EXT1/HSPG/JAK/STAT3 axis as a central regulator of tumor progression. These findings suggest that EXT1 may have value as a diagnostic marker and therapeutic target in aggressive breast cancers.

## Results

### EXT1 expression and heparan sulfate levels correlate with breast cancer aggressiveness

We aim to determine whether, in breast cancer cell lines, the extracellular HSPG landscape is subtype-dependent. By staining cell lines with the specific heparan sulfate antibody (10E4) ^23^, we found that mesenchymal cell lines (MDA-MB-231, MDA-MB-157, and Hs-578-T) exhibited significantly higher HSPG levels relative to epithelial cell lines (MDA-MB-468, ZR-75-1, and MCF-7) (Fig. 1a). To eliminate the possibility of cell line-specific effect, we stained engineered human mammary epithelial (HMLE) cells ^24^ and their mesenchymal-derived naturally arising mesenchymal cells (NAMEC) ^25^ with 10E4. We found that similar to breast cancer cell lines, NAMEC cells displayed a significantly higher HSPG level than their HMLE parental cells (Fig. 1b). To confirm the specificity of the 10E4 antibody, we treated MDA-MB-231 cells with heparinase III ^26^, an enzyme that cleaves the HS modification. This treatment resulted in a dose-response decrease in the percentage of 10E4 positive cells (Fig. S1a). Together, we identified that the HSPG landscape correlates with the mesenchymal state, leading to further determining the cellular mechanisms that differentially regulate HSPG biosynthesis in breast cancer cells.

**Fig. 1:**
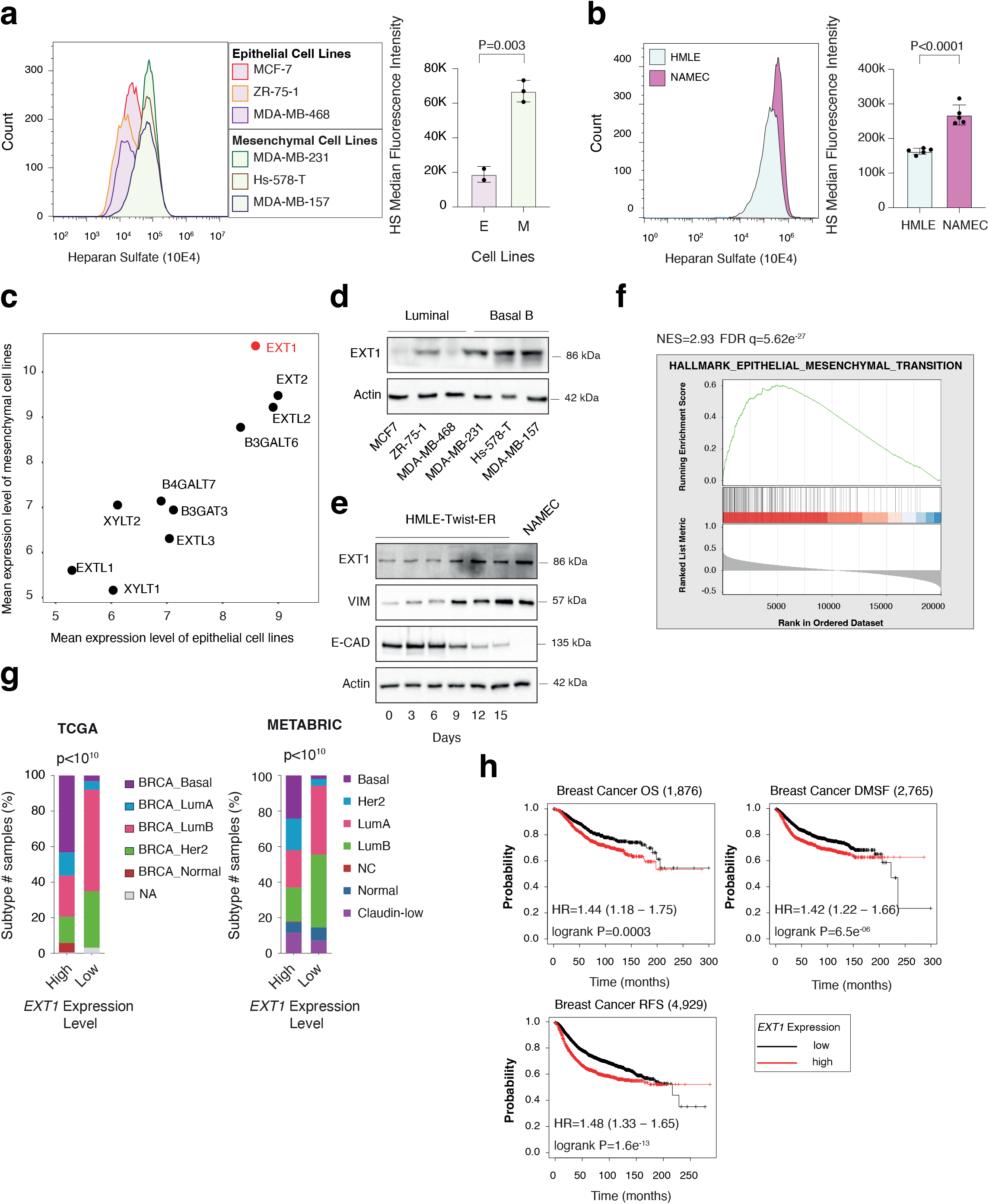
EXT1 expression is elevated in mesenchymal-like cells. **a** HSPGs levels are upregulated in mesenchymal cell lines. The indicated breast cancer-derived cell lines were subjected to FACS analysis using heparan sulfate-specific 10E4 antibody (left). The HS median fluorescence intensity was calculated for the epithelial and mesenchymal cell lines (right). The p-value was determined by Student’s *t*-test. **b** HSPGs levels are upregulated in NAMEC vs. HMLE cells. NAMEC, an HMLE-derived cell line that spontaneously acquired the mesenchymal state. The indicated HMLE and NAMEC cell lines were subjected to FACS analysis using the indicated 10E4 antibody (left). The HS median fluorescence intensity was calculated for the epithelial and mesenchymal cell lines (right). The p-value was determined by Student’s *t*-test. **c** *EXT1* demonstrated elevated expression in mesenchymal cells. Cancer cell lines were divided into epithelial (n=378 cell lines) and mesenchymal (n=150 cell lines) groups based on the expression of known mesenchymal markers. The expression of the 10 HS biosynthesis genes was compared with the mean expression in each group. **d** The EXT1 protein level is upregulated in mesenchymal breast cancer cell lines. Cells were lysed and subjected to immunoblotting using the indicated antibodies. **e** EXT1 expression is upregulated during the EMT program. HMLE-Twist-ER cells were treated with 4-hydroxytamoxifen (OHT) to induce EMT for 15 days. The cells were collected every three days, then lysed and subjected to immunoblotting using the indicated antibodies. **f** *EXT1* expression in breast cancer patients correlates with the hallmark of EMT. Breast cancer patients’ gene expression data were generated by the TCGA (PanCancer Atlas project) and analyzed using the cBioportal web tool (https://www.cbioportal.org). In these samples, the expression of *EXT1* was compared to the whole transcriptome (~20,000 genes). The genes were then ranked based on the obtained Spearman’s rank correlation coefficient followed and subjected to gene set enrichment analysis (GSEA). GSEA computed the normalized enrichment score (NES) and false discovery rate (FDR) values. **g** *EXT1* expression is elevated in the aggressive breast cancer subtypes. Breast cancer samples were divided into two groups based on high and low *EXT1* expression (one standard deviation above or below the mean). For each group, the percentage of breast cancer subtypes is color-coded. The breast cancer data was obtained from the TCGA (PanCancer Atlas project) (left) or the METABRIC (right) databases, and the p-values were calculated by the cbioportal. Lum=luminal. **h** *EXT1* expression is associated with overall survival (OE), relapse-free survival (RFS), and DM-free survival (DMFS) Kaplan-Meier survival plots for patients with breast cancer were divided into high *EXT1* expression (“high” (red)) and low (“low” (black)). The numbers in parentheses indicate the total number of patients. These plots were generated by the Kaplan-Meier plotter website. The *EXT1* (201995_at Affymetrix ID symbol) was used for all the analyses. The hazard ratio (HR) and the log-rank p-value (p) were determined by the analysis tool.

We set out to systematically identify which of the HSPG biosynthesis enzymes regulates the differential level of this proteoglycan between mesenchymal and epithelial cancer cell lines. Since the E104 recognizes the naïve form of the HS chain ^23^, we limited our analysis to 10 genes encoding for the HS synthesis initiation and chain elongation enzymes (Fig. S1b). Previously, we segregated the cancer cell lines of the MERAV database (http://merav.wi.mit.edu/) ^27^ into epithelial (378 cell lines) and mesenchymal (150 cell lines) groups according to their transcriptomes ^28^. Here, we analyzed the expression profile of the heparan sulfate biosynthesis genes (KEGG ID: hsa00534), as identified by the Kyoto encyclopedia of genes and genomes (KEGG) database (https://www.kegg.jp/pathway/map=hsa00534&keyword-Glycosaminoglycan) ^29,30^ in these two groups. As a result, we found that eight HSPG biosynthesis genes were significantly upregulated in mesenchymal cells (Fig. S1c). Furthermore, since *EXT1* exhibited the highest expression level in mesenchymal cells (Fig. 1c), we decided to characterize its contribution to the EMT program and, subsequently, to cancer cell migration.

Next, we verified this bioinformatic analysis and demonstrated that at both the mRNA (Fig. S2a) and protein levels (Fig. 1d), the EXT1 expression level was upregulated in mesenchymal relative to epithelial breast cancer cell lines. Finally, to determine whether the EMT program directly regulates EXT1 expression, we applied the HMLE-Twist-ER-based system, in which Twist, an EMT-transcription factor, is activated by hydroxytamoxifen (OHT) treatment ^31^. We found that over 15 days of treatment, these cells upregulated the expression of known mesenchymal markers such as vimentin (VIM) and cadherin 2 (*CDH2* (N-cadherin)) and suppressed the epithelial marker cadherin 1 (*CDH1* (E-cadherin)) (Fig. 1e and S2b). Similar to the known mesenchymal markers, EXT1 also exhibited a gradual increase in its mRNA and protein levels over the course of OHT treatment. Additionally, EXT1 demonstrated a higher expression level in NAMEC relative to HMLE cells, which corresponds to the HSPG profile (Fig. 1b).

We then assessed whether also, in patient-derived cancer samples, EXT1 expression correlates with mesenchymal markers and cancer aggressiveness. Hence, we determined Spearman’s rank correlation coefficient in 25 different cancer types between *EXT1* expression and the whole transcriptome (~20,000 genes), and subjected it to gene set enrichment analyses (GSEA) ^35^. This analysis revealed that *EXT1* expression significantly correlates with the hallmark of the epithelial-mesenchymal transition gene set in all examined tumor types (Fig. S2c), including breast cancer (Fig. 1f). Next, we examined whether high *EXT1* expression levels in patient-derived breast cancer samples correlate with more aggressive subtypes. To this end, we analyzed two different databases available on cBioportal (https://www.cbioportal.org) ^32^, the cancer genome atlas (TCGA, PanCancer Atlas project) ^33^ and the molecular taxonomy of breast cancer international consortium (METABRIC) ^34^. We found that in both studies, *EXT1* expression is significantly elevated in luminal B, Her2, and basal breast cancer subtypes relative to low-grade luminal A samples (Fig. S2d). Additionally, we identified that samples highly expressing EXT1 (EXT1-high) are significantly enriched with high-grade breast cancer subtypes (basal and claudin-low) in comparison to EXT1-low samples (Fig. 1g).

Finally, by analyzing the Kaplan-Meier Plotter tool (http://kmplot.com/analysis/) ^36^ and the cbioportal web tool, we identified a significant association between high *EXT1* expression levels and poor patient outcome in breast cancer overall survival (OS), relapse-free survival (RFS), and distant metastasis-free survival (DMFS) (Fig. 1h and S2d). Collectively, we demonstrated that EXT1 expression is elevated in the more aggressive breast cancer subtypes and correlates with mesenchymal markers. Thus, indicating that this enzyme can function as a regulatory component in EMT-dependent HS synthesis.

### EXT1 is sufficient to induce HSPG synthesis and the EMT program

Since we identified a direct correlation between EXT1 expression and HSPG levels, we assessed whether manipulating its expression affects HS biosynthesis. Accordingly, we ectopically expressed EXT1 in the breast cancer-derived epithelial cell lines MCF-7 and MDA-MB-468 (Fig. 2a). Then, by staining these cells with 10E4, followed by FACS analysis, we found that in both cell lines, EXT1 expression significantly elevated HSPG levels (Fig. 2b). Followingly, we systematically determined whether EXT1 expression is sufficient to induce the EMT program. Therefore, we subjected wild-type MCF-7 (VC) and EXT1 overexpressing (EXT1-FLAG) cells to comparative RNA-seq analysis (Table S1). Then the genes were ranked based on their expression ratio and applied GSEA. This analysis revealed that ectopic expression of EXT1 (Fig. S3a) significantly induces the hallmarks of the EMT gene set (Fig. 2c and Table S1). Furthermore, by qRT-PCR, we validated that EXT1 (EXT1-FLAG) is sufficient to upregulate the expression of the known mesenchymal markers *CDH2,* fibronectin (*FN1*), cluster of differentiation 44 (*CD44*), *VIM*, Snail Family Transcriptional Repressor 1 and 2 (*SNAI1* and *SNAI2*) (Fig. 2d). In contrast to EXT1-FLAG, the catalytic dead mutant (EXT1-D164A-FLAG) ^20^ failed to elevate the HSPG level (Fig. 2b) and mesenchymal gene expression (Fig. 2d). Hence, demonstrating the essentiality of EXT1 enzymatic activity to both HSPG synthesis and the execution of the EMT program.

**Fig. 2:**
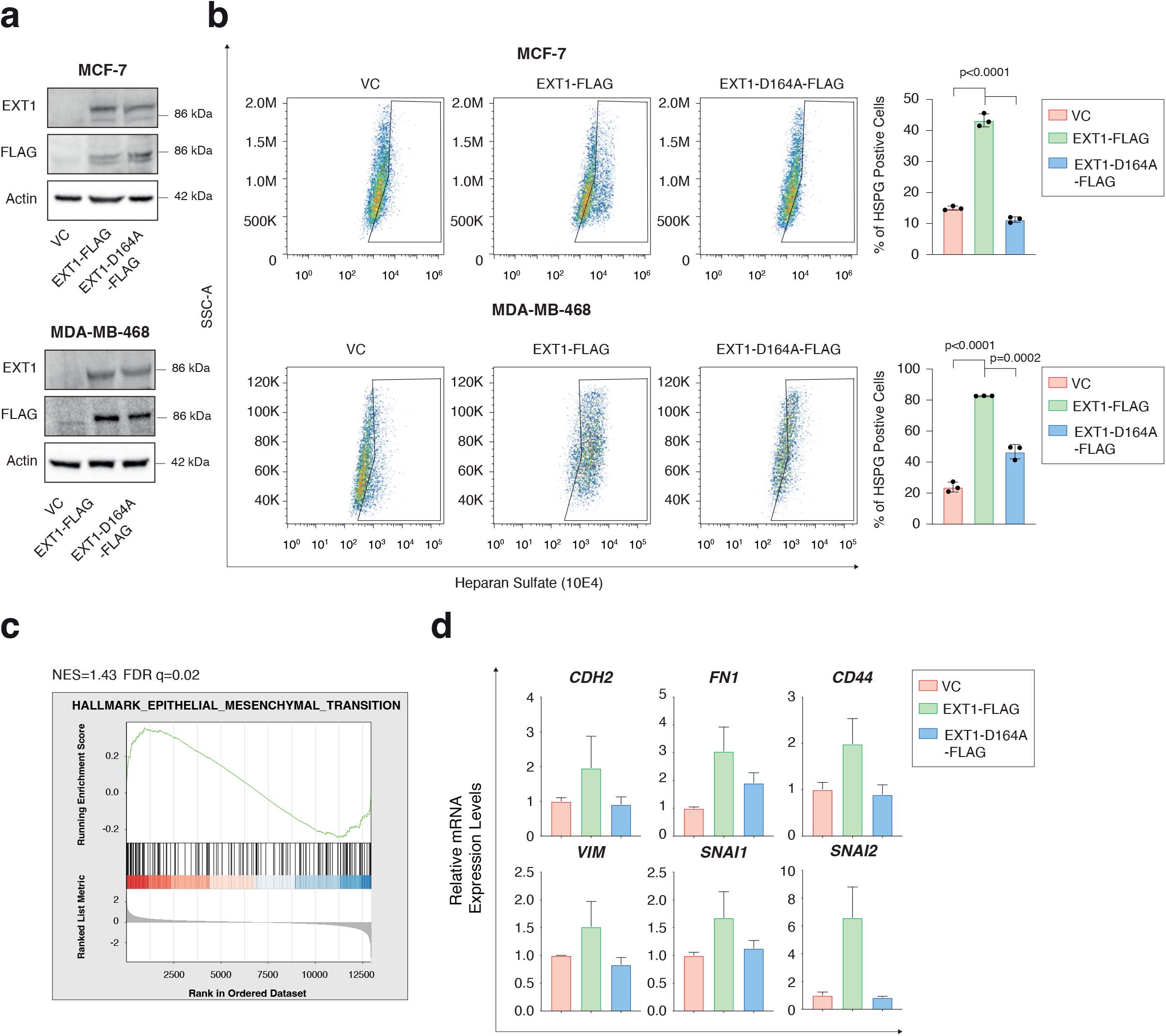
EXT1 overexpression increase heparan sulfate levels. **a** Immunoblots representing WT cells, EXT1 overexpression (EXT1-FLAG) and the catalytic dead mutant (EXT1-D164A-FLAG) MCF-7 (upper) and MDA-MB-468 (lower). Cells were lysed and subjected to immunoblotting using the indicated antibodies. VC-vector control. **b** EXT1 activity is essential for HSPG formation. The same cells as (a) were subjected to FACS analysis using the indicated 10E4 antibody (left). The HS median fluorescence intensity was calculated for each cell line (right). **c** EXT1 induces the overexpression of the mesenchymal gene set. MCF-7 cells expressing vector control (VC) WT and EXT1-FLAG were subjected to comparative RNA-Seq analysis. For each gene, the ratio between WT and EXT1-FLAG was calculated, ranked, and then subjected to gene set enrichment analysis (GSEA). GSEA computed the normalized enrichment score (NES) and false discovery rate (FDR) values. **d** Selected mesenchymal markers mRNA level is upregulated in EXT1-FLAG MCF-7 breast cancer cell lines. The relative mRNA level of selected mesenchymal markers was determined using qRT-PCR in WT cells, EXT1 overexpression (EXT1-FLAG), and the catalytic dead mutant (EXT1-D164A-FLAG) MCF-7 cells. Each value represents the mean ± SD for n=3.

### EXT1 is sufficient to promote cell migration and tumor formation

After observing that ectopic expression of EXT1 in breast cancer-derived epithelial cells elevates HSPG synthesis and subsequently induces the EMT program, we set to determine the effect on cell aggressiveness. Using the Incucyte Live-Cell analysis system, we found that ectopic expression of EXT1 in MDA-MB-468 (EXT1-FLAG) significantly enhanced the cell’s ability to heal the wound density (Fig. 3a and S3b). Furthermore, by applying Boyden chamber-based transwell migration (Fig. 3b and S3c) and invasion assay (Fig. 3c and S3d), EXT1 overexpression in both MCF-7 and MDA-MB-468 (EXT1-FLAG) resulted in increased migratory and invasive capabilities. However, overexpression of the catalytic dead form of EXT1 (EXT1-D164A-FLAG), which failed to enhance HSPG synthesis, did not change the cell characteristics. To extend the connection between HSPG formation and cell migration, we treated MDA-MB-468 cells overexpressing EXT1 (EXT1-FLAG) with heparinase III, which significantly reduces the cell’s ability to migrate (Fig. S3e). Collectively we demonstrated the EXT1-HSPG synthesis axis as a regulator of cell migration.

**Fig. 3:**
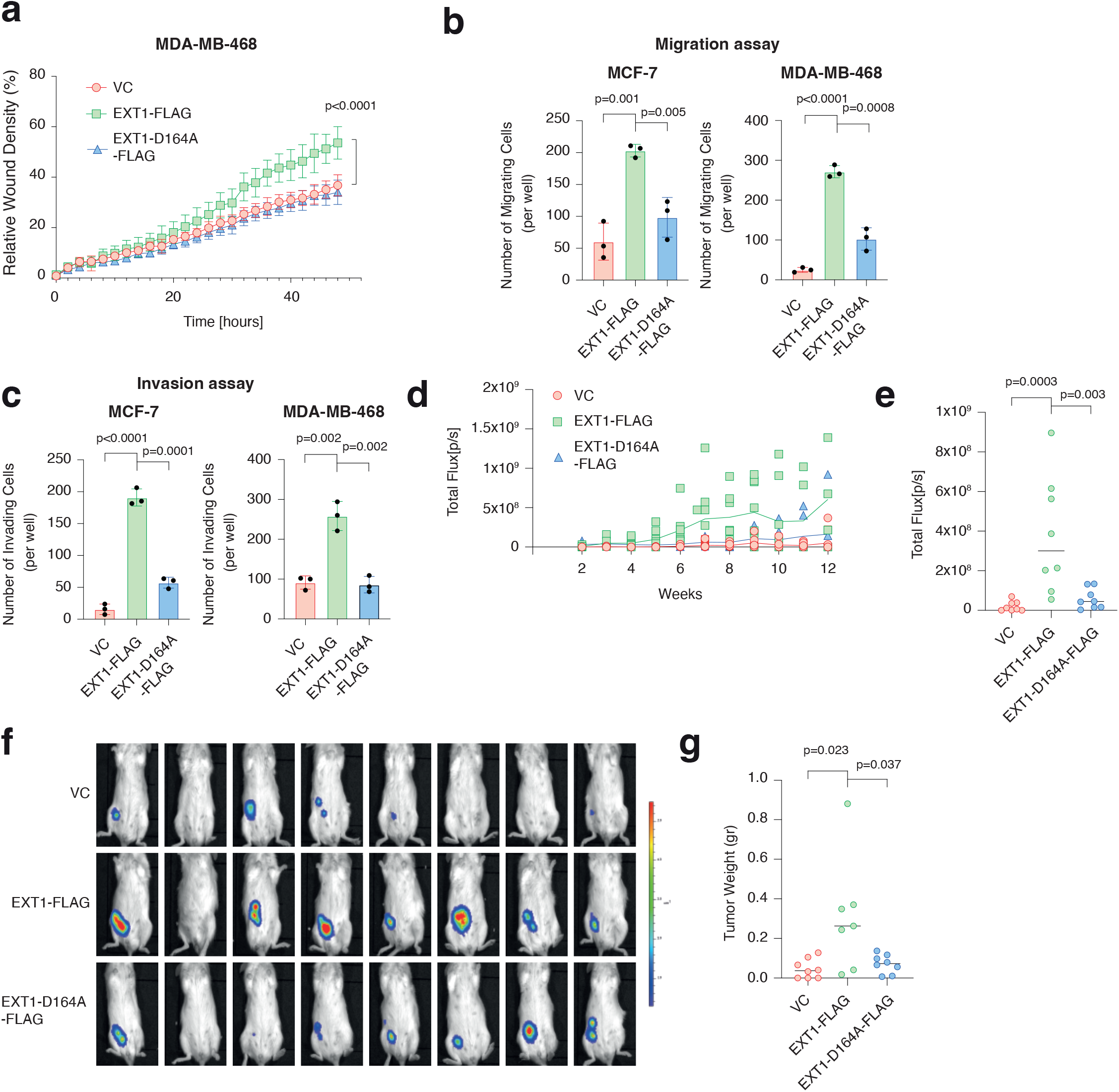
EXT1 overexpression increases the epithelial cell migratory capabilities. **a** Real-time quantification of relative wound density for the indicated WT, EXT1-FLAG, and EXT1-D164A-FLAG cells. The cells were monitored for the indicated time. Each bar represents the mean ± SD for n=8. The p-value was determined by Student’s *t*-test. **b** EXT1 overexpression induces MCF-7 and MDA-MB-468 cell migration. The migratory capability of the different samples was determined in a Boyden chamber-based transwell migration assay. Quantification of data is reported as the number of migrated cells per 25,000 seeded cells; each bar represents the mean ± SD for n=3. The p-value was determined by Student’s *t*-test. **c** EXT1 overexpression induces MCF-7 and MDA-MB-468 cell invasion. Cells were infected and treated as in (b), and the number of the matrigel-invading cells was measured. Quantification of data is reported as the number of invading cells per 20,000 seeded cells; each value represents the mean ± SD for n=3. The *p*-value was determined by Student’s *t*-test. **d** Kinetics of EXT1 expression effect on tumor growth was determined by the bioluminescence machine. *In vivo* tumor growth kinetics of WT MDA-MB-468 (VC), EXT1-FLAG, and EXT1-D164A-FLAG overexpressing cells were determined every two weeks in NOD-SCID mice. **e** Quantification of the bioluminescence images. Each ball represents the bioluminescence in a mouse n=8. The p-value was determined by the Mann-Whitney U test. **f** Representative images of NOD-SCID mice after 12 weeks post-injection with MDA-MB-468 cells expressing VC, EXT-FLAG, or EXT1-D164A-FLAG were captured using bioluminescence. The color bar represents the intensity of luminescence. **g** The tumors were harvested from the mice, weighed, and presented as a graph. The p-value was determined by Student’s *t*-test.

Next, we assessed the role of EXT1 in tumor formation *in vivo.* Accordingly, we injected luciferase-expressing plasmid and GFP-labeled MDA-MB-468 cells (VC, EXT1-FLAG, or EXT1-D164A-FLAG) into the mammary fat pad of female NOD-SCID mice. By real-time monitoring of luciferase activity ^37^ for 12 weeks, we found that EXT1 expression significantly induced tumor growth kinetics (Fig. 3d) and resulted in larger tumor size (Fig. 3e and 3f) and weight (Fig. 3g and S3f). Collectively, these results highlight that the ectopic expression of enzymatically active EXT1 in epithelial cells is sufficient to elevate HSPG formation, subsequently increasing their migratory capabilities.

### EXT1 is a potent regulator of breast cancer aggressiveness

After determining that ectopic expression EXT1 is sufficient to induce HSPG formation and the EMT program, we examined whether its silencing will attenuate the cells’ aggressive characteristics. First, by applying the CRISPR-Cas9 system, we knockout EXT1 in the basal B breast cancer cell line, MDA-MB-231 (EXT1-KO) (Fig. 4a), which significantly reduced HSPG levels (Fig. 4b). Then, to avoid any off-target effect, we ectopically expressed the WT EXT1 (EXT1-FLAG) or the catalytic dead mutant (EXT1-D164A-FLAG) in the background of EXT1 knockout. We found that, as opposed to the ectopic expression of the WT form, which resulted in a significant elevation in HSPG levels, the catalytic dead mutant failed to induce this proteoglycan synthesis (Fig. 4b).

**Fig. 4:**
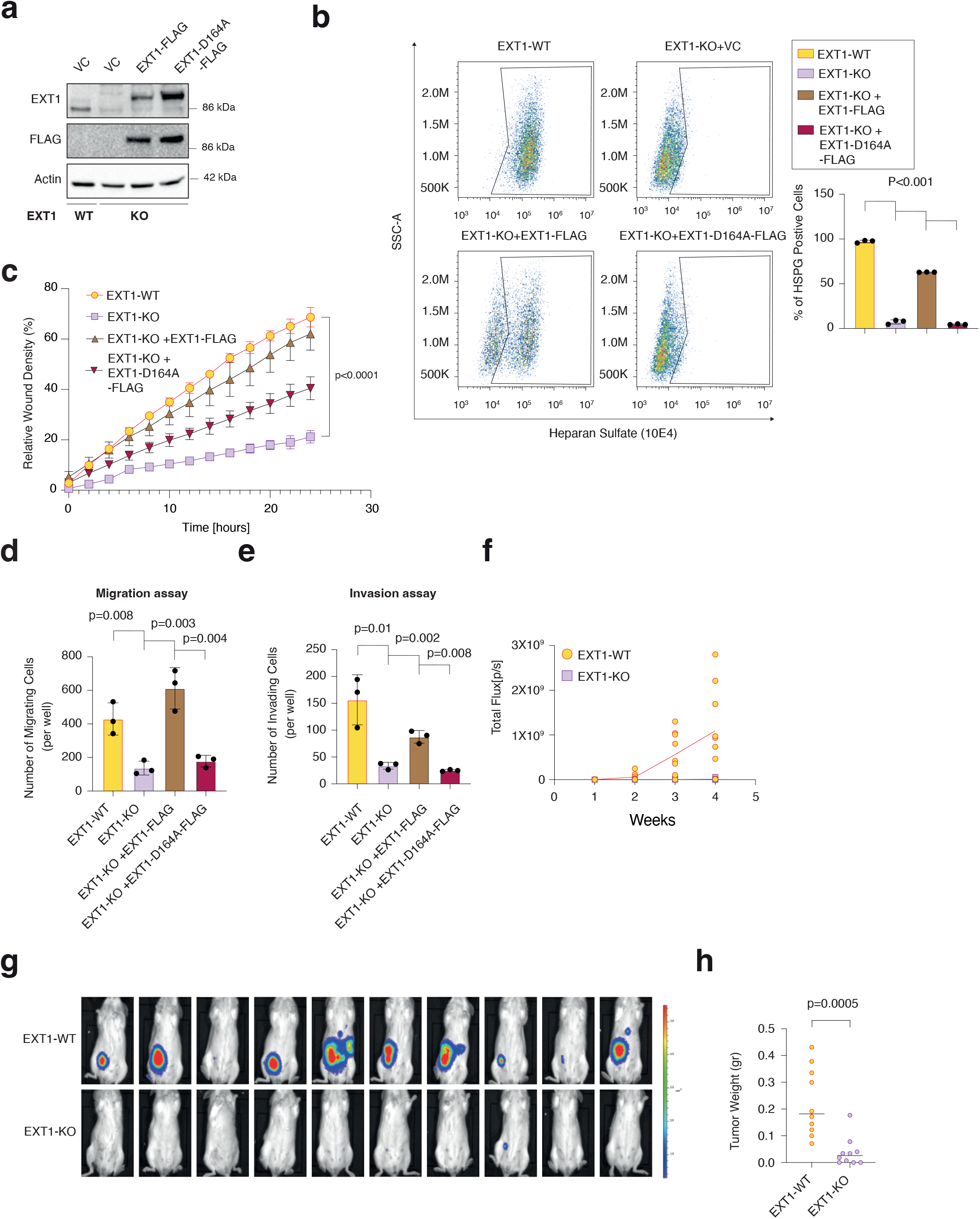
EXT1 expression regulates breast cancer aggressiveness. **a** Immunoblot representing EXT1 knockout (EXT1-KO) in MDA-MB-231. EXT1 knockout was generated using the CRISPR-Cas9 system, followed by separating the cells into single clones. FLAG-tagged EXT1 (EXT1-FLAG) and catalytic dead mutant (EXT1-D164A-FLAG) were reintroduced into EXT1-KO cells. Cells were lysed and subjected to immunoblotting using the indicated antibodies. **b** EXT1 activity is essential for HSPG formation. The same cells as (a) were subjected to FACS analysis using the indicated 10E4 antibody (left). The HS median fluorescence intensity was calculated for each of the samples (right). Each bar represents the mean ± SD for n=3. The p-value was determined by Student’s *t*-test. **c** The same cells as (a) were subjected to real-time quantification of relative wound density. The cells were monitored for the indicated time. Each bar represents the mean ± SD for n=8. The p-value was determined by Student’s *t*-test. **d** EXT1 loss inhibits cell migration in breast cancer cells. The same cells as (a) were subjected to Boyden chamber-based transwell migration assay. Quantification of data is reported as the number of migrated cells per 20,000 seeded cells; each bar represents the mean ± SD for n=3. The p-value was determined by Student’s *t*-test. **e** The same cells as (a) were subjected to matrigel-based invasion assay. Quantification of data is reported as the number of migrated cells per 25,000 seeded cells; each value represents the mean ± SD for n=3. The *p*-value was determined by Student’s *t*-test. **f** The kinetics effect of EXT1 loss on tumor growth was determined by the bioluminescence machine. *In vivo* tumor growth kinetics of WT MDA-MB-231 (EXT1-WT), and EXT1-KO cells were determined every two weeks in NOD-SCID mice. **g** Representative images of mice after 4 weeks postinjection. The images were captured using bioluminescence. The color bar represents the intensity of luminescence. **h** The tumors were harvested from the mice, weighed, and presented as a graph. The p-value was determined by Student’s *t*-test.

To determine the correlation between HSPG reduction and cell migration, we applied the Incucyte Live-Cell analysis system. We found that EXT1-KO cells exhibit a significant decrease in wound healing kinetics relative to WT (EXT1-WT) cells (Fig. 4c and S4a). Additionally, the loss of EXT1 (EXT1-KO) significantly affected MDA-MB-231 ability to migrate (Fig. 4d and S4b) and invade (Fig. 4e and S4c). We verify that the specificity of these effects was due to EXT1 loss and subsequent HSPG reduction, by overexpressing, in the knockout background, the WT (EXT1-KO+EXT1-FLAG) and the catalytic dead mutant (EXT1-KO+EXT1-D164A-FLAG) forms. We found that in contrast to the WT form, which rescued the KO effect, the catalytic dead mutant failed to restore the cell’s ability to migrate and invade, thus indicating that EXT1 enzymatic activity is essential for cell migration. Moreover, treating WT cells with heparinase III also significantly reduced the cell’s ability to migrate (Fig. S4d). Finally, to eliminate a cell line-specific effect, we knocked out EXT1 from Hs-578-T (Fig. S4e), another basal B breast cancer cell line. We found that EXT1 loss in this cell line (EXT1-KO) reduces the expression of known EMT markers such as *CDH2,* Zinc Finger E-Box Binding Homeobox 1 (*ZEB1), CD44, FN1,* Twist Family BHLH Transcription Factor 1 (*TWIST1*), *SNAI2,* and interleukin 6 (*IL-6*) (Fig. S4f). Moreover, EXT1 loss in Hs-578-T (EXT1-KO) significantly reduces the ability of the cells to migrate (Fig. S4g). Together, these results further support the essential role of EXT1 as a regulator of cancer cell aggressiveness through its function in HSPG synthesis.

We then determine the role of EXT1 in tumor formation in mice. To this end, we injected luciferase-expressing plasmid and GFP-labeled MDA-MB-231 cells (WT and EXT1-KO) into the mammary fat pad of female NOD-SCID mice. Then, we real-time monitored the tumor growth kinetics for four weeks using the noninvasive bioluminescence imaging system. We found that in these mice, the growth kinetics of EXT1-KO cells was significantly lower than WT cells (Fig. 4f and 4g). Additionally, the average weight of tumors generated from EXT1-KO cells was significantly reduced than those from WT cells (Fig. 4h and S4h). Collectively, these findings underline the role of EXT1 expression and activity in cell migration and tumor growth kinetics.

### EXT1 as a regulator of STAT3 signaling

To identify the cellular mechanisms by which EXT1 governs breast cancer aggressiveness, we submitted wild-type (EXT1-WT) and knockout (EXT1-KO) MDA-MB-231 cells to a comparative CEL-Seq analysis ^38^ (Table S2). Then, we ranked the genes based on their expression ratio followed by GSEA to systematically characterize the gene sets downregulated upon EXT1 loss. This analysis identified a significant reduction in 15 hallmarks (Fig. S5a), including the “Hallmark of IL6 STAT3 Signaling,” which has the lowest normalized enrichment score (NES, −1.78) (Fig. 5a) and “Hallmark of EMT” (NES, −1.48) (Fig. S5b). Hence, EXT1 loss induces significant changes in the cell’s transcriptome through the repression of signaling pathways such as JAK/STAT3.

**Fig. 5:**
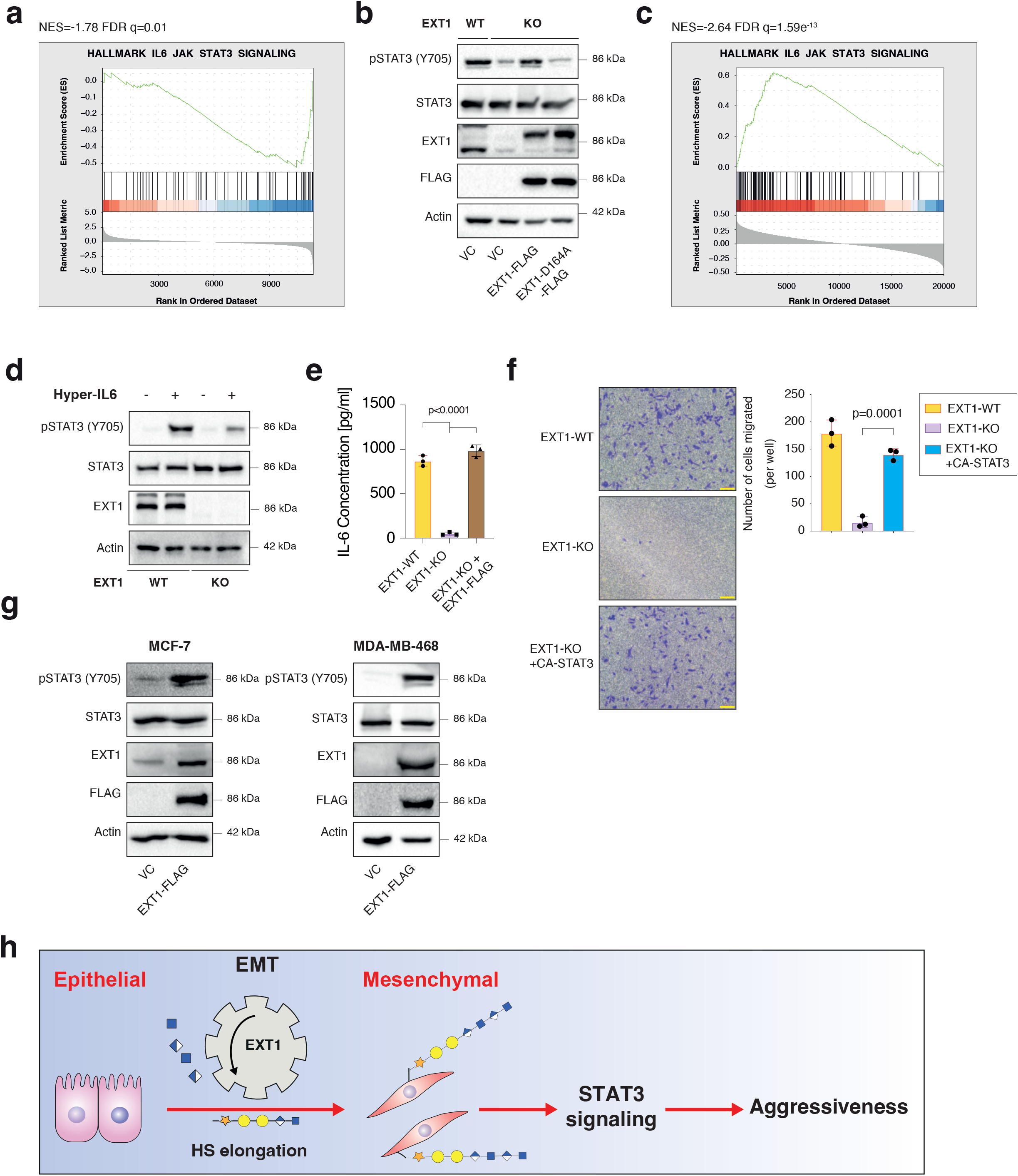
EXT1 loss impairs STAT3 signaling. **a** EXT1 loss leads to a reduction in the gene expression pattern for “hallmark IL-6/JAK/STAT3 signaling.” MDA-MB-231 WT cells and EXT1-KO-1 were subjected to CEL-Seq analysis. The expression ratio of all genes was calculated and ranked based on the relative expression in EXT1-WT and EXT1-KO. The samples were subjected to GSEA. The FDR q-value was computed by GSEA. **b** STAT3 phosphorylation correlates with EXT1 expression. Immunoblots representing WT MDA-MB-231 expressing VC, EXT1 knockout (EXT1-KO), and overexpression of the full-length variant (EXT1-FLAG) or the catalytic dead mutant (EXT1-D164A-FLAG) in the KO background. Cells were lysed and subjected to immunoblotting using the indicated antibodies. **c** EXT1 expression in breast cancer patients correlates with the hallmark of IL6/JAK/ STAT3 signaling. Breast cancer patients’ gene expression data were generated by the TCGA (PanCancer Atlas project) and analyzed using the cBioportal web tool (https://www.cbioportal.org). In these samples, the expression of EXT1 was compared to the whole transcriptome (~20,000 genes). The genes were then ranked based on the obtained Spearman’s rank correlation coefficient and then subjected to GSEA. GSEA computed the normalized enrichment score (NES) and false discovery rate (FDR) values. **d** Loss of EXT1 expression results in STAT3 signaling inhibition. EXT1-WT and EXT1-KO cells were starved with 0% FBS medium for 16 h and treated with 0 and 50 μl of media from HEK-293 cells generating Hyper IL-6 for one hour. Cells were subjected to immunoblot using the indicated antibodies. **e** IL-6 level is reduced in EXT1-KO cells growth media. After 36h, cell growth media was collected from each of the indicated samples and the IL-6 level was measured using a specific ELISA kit (n = 3). The p-value was determined by Student’s *t*-test. **f** Constitutively activated STAT3 (CA-STAT3) improves the migration of EXT1-KO cells. The migratory capability of the different samples was determined using Boyden chamber-based transwell assay. Representative images of each sample (left). Scale bar=100μm. Quantification of data is reported as the number of migrated cells per 20,000 seeded cells (right). Each bar represents the mean ± SD for n=3. The p-value was determined by Student’s *t*-test. **g** EXT1 overexpression in epithelial MCF-7 and MDA-MB-468 cell lines induces STAT3 signaling. Cells expressing vector control (VC), and EXT1-FLAG, were lysed and subjected to immunoblotting using the indicated antibodies. **h** A schematic representation of EXT1’s role in JAK/STAT3-mediated cell aggressiveness.

We validated the CEL-seq analysis to establish the EXT1/HSPG/JAK/STAT3 axis. We found a direct correlation between EXT1 expression and STAT3 phosphorylation (pSTAT3) on tyrosine 705 (Y705). Specifically, EXT1 loss in MDA-MB-231 cells (EXT1-KO) is sufficient to reduce the pSTAT3 levels (Fig. 5b), whereas ectopic expression of the full-length EXT1 in the knockout background (EXT1-KO+EXT1-FLAG), elevated back pSTAT3 levels. Furthermore, in contrast to the WT form, the catalytic dead mutant (EXT1-KO+EXT1-D164A-FLAG) failed to elevate pSTAT3 levels, emphasizing the essentiality of EXT1 activity for STAT3 activation. To further validate this axis, we treated MDA-MB-231 cells with heparinase III, which reduced the levels of both HSPG (Fig. S1a) and pSTAT3 (Fig. S5c), linking HSPG formation with JAK/STAT3 activation. Finally, in breast cancer patients, EXT1 expression significantly correlated with IL-6/JAK/STAT3 signaling (Fig. 5c). These findings indicate that EXT1 functions as an upstream regulator of the JAK1-STAT3 signaling cascade.

To further unravel EXT1 as a regulator of the JAK/STAT3 cascade, we treated both WT and EXT1-KO cells with Hyper-IL6 ^39^, a fusion protein combining IL-6 and soluble IL6-receptor (sIL6R), which is a potent inducer of STAT3 activation ^13^. We found that Hyper-IL6 substantially induced pSTAT3 in WT cells, whereas this phosphorylation was attenuated in EXT1-KO cells (Fig. 5d). Next, we applied the STAT3 downstream targets as a readout for JAK/STAT3 activation. Among the STAT3 target genes that were significantly downregulated upon EXT1 loss in our CEL-Seq analysis (Table S2) was IL-6 (Fig. S5d). This cytokine is a known STAT3 target ^40^ that regulates cancer aggressiveness and the EMT program ^41–43^. Therefore, we verified that IL-6 mRNA (Fig. S5e) and secreted protein levels (Fig. 5e) are downregulated upon EXT1 loss. However, ectopic expression of EXT1 in the background of KO cells (EXT1-KO-+EXT1-FLAG) elevated IL-6 to a similar level detected in the WT cells (Fig. 5e). Finally, to further determine that STAT3 signaling mediated EXT1’s effect on cell migration, we over-expressed the constitutively activated STAT3 (A662C, N664C, V667L, (CA-STAT3)) ^44^ in EXT1-KO cells. We found that CA-STAT3 significantly increased IL-6 mRNA (Fig. S5e) level and the number of migratory cells compared to EXT1-KO cells (Fig. 5f). Collectively, these results demonstrate that the EXT1 regulates cell migration through the HSPG/JAK1/STAT3 axis.

To further establish the EXT1/HSPG/JAK/STAT3 axis, we overexpressed EXT1 in the epithelial cell lines MCF-7 and MDA-MB-468, which induced HSPG synthesis (Fig. 2b) and was sufficient to increase pSTAT3 (Fig. 5g). Additionally, to verify that EXT1-dependent pSTAT3 is mediated by JAK1/2, we treated EXT1 overexpression MDA-MB-468 cells (EXT1-FLAG) with ruxolitinib (RUXO), a potent inhibitor of JAK1/2^45^, which abolished the STAT3 phosphorylation (Fig. S5f). Together these results demonstrated that EXT1 governs cancer cell migration through its essential role in the HSPG/JAK/STAT3 axis.

## Discussion

We identified that EXT1, through its role in HSPG synthesis, is a central regulator of breast cancer aggressiveness. By analyzing patient-derived databases, we demonstrated that EXT1 expression is upregulated in high-grade breast cancer subtypes and was associated with poor prognosis. Moreover, we showed that EXT1 expression is induced by the EMT program. To establish EXT1 essential role in breast cancer aggressiveness, we manipulated its expression level and determined the effects on the cell’s physiology. Specifically, we showed that ectopic expression of EXT1 in epithelial cell lines induced HSPG synthesis and, subsequently, their capability to migrate, invade, and initiate tumors in mice. In contrast, EXT1 knockout in the highly aggressive MDA-MB-231 mesenchymal-like breast cancer cell line reduces HSPG level, migratory and invasion capabilities, and tumor initiation abilities. Additionally, EXT1 loss inhibited the JAK/STAT3 signaling cascade, a central regulator of cancer cell aggressiveness (Fig. 5h). These findings indicate a novel EXT1/HSPG/JAK/STAT3 axis that regulates cellular migration and thus provides a direct link between metabolic enzymes synthesizing proteoglycans and oncogenic signaling pathways.

Syndecans and glypicans are the major families of cell surface proteins that are modified by HS ^14^ The syndecan family is composed of four members (SDC1-4) ^7^, where syndecan-1 (SDC1, (CD138)) is expressed in various types of tissues and tumors ^46^. In breast cancers, SCD1 has been implicated in promoting tumor growth, invasion, metastasis, and chemoresistance ^11,47^ and is significantly associated with poor patient prognosis ^48^. Moreover, the same study showed that SCD1 silencing attenuated STAT3 signaling ^12^. Thus, we speculate that most of the EXT1/HSPG-dependent cellular effects we observed in our study might be mediated by SDC1.

The HS synthesis machinery includes both EXT1 and EXT2, which can form a hetero-oligomers complex. This complex retains significantly higher glycosyltransferase and polymerizing activities than each enzyme by itself ^18^. Despite their similarity, several studies revealed physiological differences between EXT1 and EXT2. For example, *in vitro* experiments demonstrated that both enzymes cannot substitute for each other ^18^. In addition, Ext1 KO mice are lethal around the time of gastrulation, whereas Ext2-null mice develop normally until embryonic day 6.0 ^21^. Furthermore, the human gene mutation database (HGMD) analysis indicates that EXT1 has higher causative alterations than EXT2 ^22^. Finally, EXT1 mutation with multiple osteochondromas is more prone to display malignant transformation than patients with EXT2 mutations ^49,50^. In this study, by applying genetic-based manipulation techniques, we demonstrate a clear correlation between EXT1 and cancer cell aggressiveness, which was inert for endogenous EXT2 expression. This failure of endogenous EXT2 to compensate for the EXT1-dependent effects on the cell’s characteristics could be due to their different, non-redundant features.

Gene expression profile studies identified EXT1 as an indicator for the NOTCH pathway activation in acute lymphoblastic leukemia (ALL) ^51^ and as a marker for disease-free survival in hepatocellular carcinoma ^52^. In breast cancer patients, upregulation of *EXT1* mRNA levels was detected in estrogen receptor (ER)-negative cells ^53^ and was suggested to function as a high-risk predictor for metastasis ^54^. Additionally, an association between EXT1 expression and cancer stemness was found in MCF-7 cells selected for doxorubicin-resistant (MCF-7/ADR) ^55^. Specifically, these selected resistant cells demonstrated cancer stem cell properties promoted by EXT1 expression. Collectively, several studies identify EXT1 as a marker for breast tumor aggressiveness, although the means by which this enzyme governs cell aggressiveness was vague. Here we introduce the novel EXT1/HSPG/JAK/STAT3 axis that provides the mechanism by which EXT1 shifts breast cancer cells to a more aggressive state.

Previously, by conducting a gene expression analysis followed by a FACS-based screen, we identified a set of metabolic genes, designated as the “mesenchymal metabolic signature” (MMS), that function as potential regulators of the EMT program ^28^. Among them are dihydropyrimidine dehydrogenase (DPYD) and glutathione peroxidase 8 (GPX8), which we established as EMT regulators^28,41^. Interestingly, this screen also identified the EXT1 that we confirmed here to mediate the aggressive characteristics induced by the EMT program. Our results reinforce the FACS-based screen results and suggest that other hits may also function as essential components of the EMT program.

HSPGs are composed of highly sulfated repeating disaccharide units ^56^ found on the cell surface and in the extracellular matrix ^5^. Due to their negative charge, the sulfated chains act as co-receptors, enhancing the interactions between ligand and their respective receptors ^57^. This ligand-receptor interaction triggers the activation of the downstream signaling pathways, such as MAPK, AKT, Wnt, and JAK/STAT3 ^7,57^. Previous studies demonstrated that a reduction in EXT1 expression levels results in shorter HS chains ^58^. This reduced HS chain length, caused by EXT1 loss, was reported to attenuate intracellular signaling, such as Wnt, in multiple myeloma cells ^26^. In this study, we explored the cellular mechanisms by which EXT1 promotes breast cancer aggressiveness. By applying unbiased CEL-Seq analysis followed by biochemical experiments, we identified the JAK/STAT3 as the major signaling pathway downregulated upon EXT1 loss. However, we cannot rule out that EXT1, through HSPG synthesis, also regulates other known signal transduction pathways. We chose to focus on JAK/STAT3 signaling cascade since it is a major regulator of cancer cell aggressiveness ^41^ and migration ^13^.

Here we demonstrate that HSPG formation is crucial in breast cancer aggressiveness. The cellular machinery synthesizing this PG involves a series of sequential reactions ^56^ that include more than 20 different enzymes ^14^. Therefore, identifying putative targets in this metabolic pathway could have a therapeutic outcome. Since we revealed that ectopic expression of EXT1 is sufficient to induce HSPG formation, we suggest this enzyme as the regulatory factor in this process. Further studies are needed to fully understand the mechanisms by which EXT1 and heparan sulfate contribute to cancer aggressiveness and determine their translational capabilities. Overall, this study contributes to the significance of the mesenchymal state in breast cancer aggressiveness and the potential of targeting this state for developing effective therapies.

## Materials and methods

### Cell lines and cell culture

The cell lines ZR-75-1, MCF-7, MDA-MB-468, MDA-MB-231, Hs-578-T, and MDA-MB-157 were obtained from ATCC and were cultured in DMEM supplemented with 10% heat-inactivated HI-FBS (Biological Industries). The immortalized human mammary epithelial cells expressing OHT-inducible Twist (HMLE-Twist-ER) and Naturally Arising MEsenchymal Cells (NAMECs) were maintained in MEGM (Lonza) growth medium. All cells were cultured at 37°C with 5% CO_2_. For EMT induction, HMLE-Twist-ER cells were treated with 4-hydroxytamoxifen (OHT) (Sigma-Aldrich, H7904) at a final concentration of 10 nM for the indicated number of days.

### Antibodies

Antibodies were obtained from the following sources: EXT-1 (sc-515144) from Santa Cruz Biotechnology, Heparan Sulfate 10E4 Antibody from Amsbio, CDH1 (3195), CDH2 (13116), β-actin (4970)\(3700), FLAG (8146), VIM (5741), p-STAT3-Tyr705 (9145), STAT3 (9139), from Cell Signaling Technology; FITC-labeled anti-HS (555427) from BD Bioscience. HRP-labeled Goat anti-mouse (115-035-003) and HRP-labeled Goat anti-rabbit (111-035-144) secondary antibodies were obtained from Jackson ImmunoResearch. Secondary antibody for FACS experiments: Donkey Anti-Mouse −488 (ab96875) from Abcam. Measurement of IL-6 levels was performed by sandwich ELISA using the human IL-6 mini ABTS ELISA development kit (Peprotech-catalog no 900-M16).

### FACS Analysis

For FACS analysis, 300,000 cells were seeded in 6-well tissue culture plates with 10% HI-FBS DMEM. The next day cells were detached using detachment buffer (0.05% EDTA in PBS) for 5 minutes and harvested in 10% HI-FBS DMEM and fixed with 4% paraformaldehyde (PFA, no-15710, Electron microscopy sciences, USA) in PBS for 10 min at room temperature. Followingly, cells were, incubated with anti-Heparan Sulfate (10E4 epitope) antibody (1:200 in PBS) for 45 minutes in ice, followed by secondary antibodies (Alexa Fluor™ 488, diluted 1:500 in PBS) incubation for 30 minutes on ice in the dark. Finally, the cells were filtered using Micron Nylon Filter Mesh Fabrics and acquired on a BD Accuri C6, and analyzed using the FlowJo software (Tree Star, Ashland, OR).

### Cell lysis and immunoblotting

Cells were rinsed once with ice-cold PBS and lysed with radioimmunoprecipitation assay (RIPA) lysis buffer (20 mM Tris [pH 7.4]), 137 mM NaCl, 10% glycerol (vol/vol), 1% Triton X-100, 0.5% (wt/vol) deoxycholate, 0.1% (wt/vol) SDS, 2.0 mM ethylenediaminetetraacetic acid (EDTA) (pH 8.0), and one tablet of EDTA-free protease inhibitor (Roche) and phosphatase inhibitor mixture mixes A and B (100X) (Bimake). The lysates were cleared by centrifugation at 13,000 rpm at 4°C in a microcentrifuge for 10 min. The protein concentration was determined by Bradford (BioRad). Proteins were denatured by adding SDS sample buffer (5x) and boiling for 5 minutes, then resolved by 10% SDS-PAGE, transferred onto a 0.45 μm polyvinylidene difluoride (PVDF) membrane (Merck), and probed with the appropriate antibodies. The images were quantified using Image Lab version 6.1.0 (BioRad).

### Analysis of breast cancer data in cBioportal

The cBioPortal for Cancer Genomics is an open-access database providing visualization and analysis tools for large-scale cancer genomics data sets (http://cbioportal.org). For gene correlation analysis, we queried EXT1 in breast invasive carcinoma (TCGA, PanCancer Atlas project) or the METABRIC (containing 1,084 or 2,509 samples, respectively). Then, we subjected the genes to co-expression analysis and downloaded the correlation plots. For gene set enrichment analysis (GSEA), Spearman’s rank correlation coefficient between the gene of interest and the whole genome was computed, downloaded, and subjected to GSEA analysis and visualization of the result, using the R package clusterProfiler ^59^ and enrichplot. For the different analyses, we selected the h.all.v7.2.symbole.gmt (Hallmarks) or C2.cp.kegg.v7.2.symbols.gmt (Curated) gene set databases.

### Cancer sample analysis

KM analyses of the breast cancer samples were analyzed and generated by the KM Plotter website (http://kmplot.com/analysis/) ^60^. Search entries: EXT1as the gene symbol (Affymetrix ID: 201995_at), the auto-select best cut-off for “split patients by” The obtained KM plots and the statistics were generated by the website.

### Virus production

HEK-293T cells were co-transfected with the pLentiCRISPR sgRNA, VSV-G envelope plasmid, and Δvpr lentiviral plasmid using X-TremeGene 9 Transfection Reagent. The supernatant containing the virus was collected 48 hours after transfection and spun for 5 min at 400g in order to eliminate cells and debris or filtered with a 0.45μm syringe filter (Lifegene, Mevo Horon, Israel).

### CRISPR/Cas9-Mediated Knockout Cell lines

We used CRISPR-Cas9-mediated genome editing to achieve gene knockout, using pLentiCRISPR v1 (Addgene Plasmid #70662) in which the sgRNA and Cas9 are delivered on a single plasmid. Editing the EXT1 locus in MDA-MB-231 cells was accomplished by infecting cells with the “pLentiCRISPR” plasmid into which a sgRNA targeting the EXT1 locus had been cloned. Cells were then subjected to single-cell cloning by limiting dilution in 96-well plates. Editing of the EXT1 locus was confirmed by assessing protein levels by immunoblot. Primers used for cloning non-targeting control (NTC), NTC-sgRNA 5’ caccgGCGCTTCCGCGGCCCGTTCAA and 5’ aaacTTGAACGGGCCGCGGAAGCGg; 3’; EXT1-gRNA1 5’ caccgGACCCAAGCCTGCGACCACG 3’ and 5’ aaacCGTGGTCGCAGGCTTGGGTCg 3’; EXT1-sgRNA2 5’ caccgGTCTGGTTCCTCGTGGTCGC 3’ and 5’ aaacGCGACCACGAGGAACCAGACg 3’; the EXT1-gRNA were designed based on Ren et al. ^26^.

### RNA preparation and qRT-PCR

Total RNA was isolated from cells using the NucleoSpin® RNA Kit (MACHEREY-NAGEL, Germany), and reverse transcription was performed using the qScript cDNA Synthesis Kit (Quantbio, USA). The resulting cDNA was diluted in DNase-free water (1:10) before quantification by real-time quantitative PCR (qRT-PCR). The mRNA transcription levels were measured using 2x qPCRBIO SyGreen, Blue Mix Hi-ROX (PCR Biosystems), and StepOnePlus (Applied Biosystems). All data were expressed as the ratio between the expression level of the target gene mRNA and actin (ß-Actin). The primers used for the qRT-PCR analysis were obtained from Integrated DNA Technology with the following sequences (Table S3):

**Table S3:**
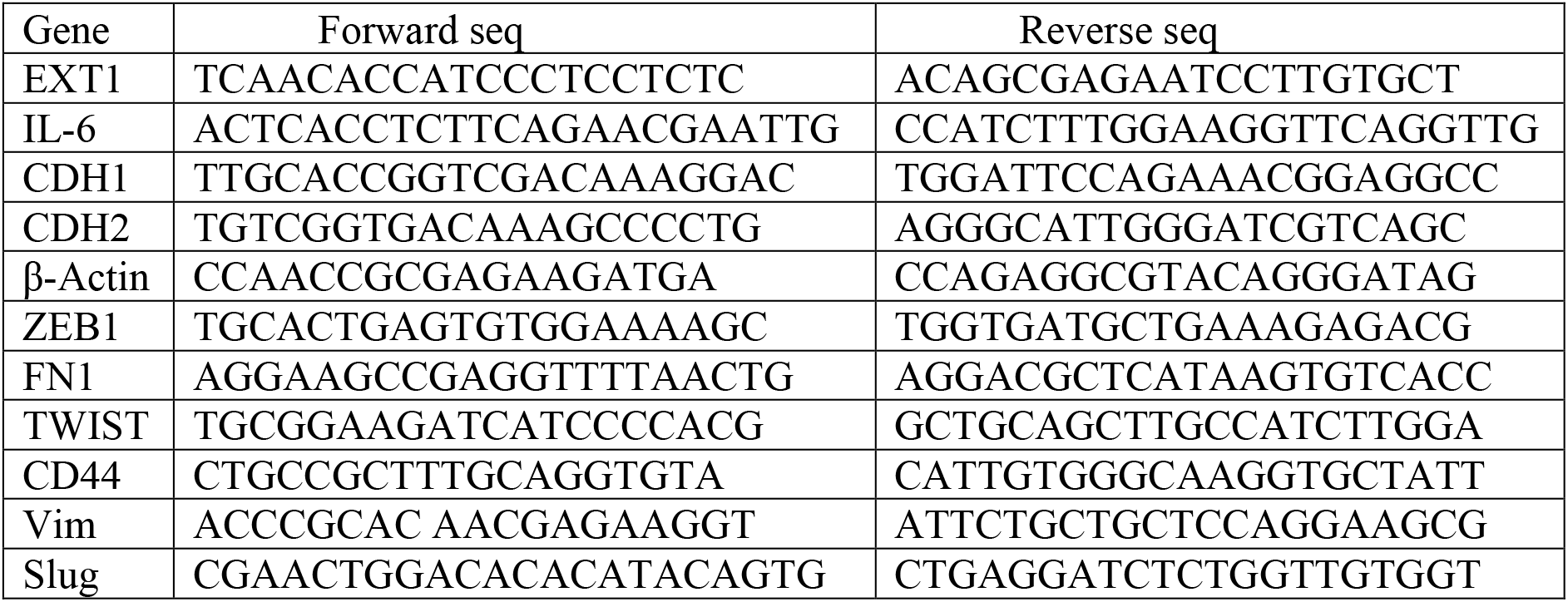
list of primers and their corresponding forward and revers sequences used for qRT-PCR.

### Transwell migration and invasion assays

Migration and invasion assays were performed using the transwell migration chamber (Costar, USA). For the migration assay, transwell inserts were rehydrated with serum-free DMEM for 30min. Then, MDA-MB-468 (2.5 x 10^3^), MCF-7 (2.5 x 10^3^), and MDA-MB-231 (2 x 10^4^) cells were suspended in 350μl of serum-free DMEM and seeded in the upper chamber. For the invasion assay, the Millicell cell culture inserts (Millipore) were coated with ECL-cell attachment matrix (08-110, Millipore) (1μg/ml) with serum-free DMEM for 1 hour. Then MDA-MB-468 (2 × 10^4^), MCF-7 (2 × 10^4^), and MDA-MB-231 (2.5 x 10^3^) cells, in serum-free DMEM, were added to the upper chamber. For both assays, to the lower chamber of the 24-well plate, 700μl of 10% HI-FBS in DMEM was added. The medium was discarded after 24 h. Cells that remained in the upper chamber were removed with cotton-tipped swabs, and the lower surface of the insert was stained with 0.5% crystal fast violet (548-62-9, Acros organics, USA). The cells were counted and captured under a Nikon Eclipse 80i microscope at 10× magnification.

### Wound Healing Assay

MDA-MB-231 and MDA-MB-468 cells (both 4 ×10^4^ cells/ well) were plated onto IncuCyte ImageLock 96-well cell culture microplates (Corning). After 18 h, the cell monolayer was scraped using a wound-maker mechanical device (Essen BioScience), washed with PBS, and examined under an inverted microscope. The wound area was monitored using the IncuCyte live-cell imaging system. Wound healing assay results were compiled from eight wells with one scratch in each well and were conducted, dependent on the cell lines, for 24h or 48h (MDA-MB-231 cells and MDA-MB-468, correspondingly).

### Heparinase III supplementation

Cells were seeded in 6cm plates, and the next day, prior to experiments the cells were washed with PBS twice and treated with Heparinase III (10mU/ml) (6145-GH-010, Amsbio) diluted in DMEM or PBS for 1hr at 37°C. After incubation, the cells are washed with PBS twice and subjected to respective experiments.

### Animal Studies

MDA-MB-231 WT GFP^+^ Luci^+^, EXT1-KO GFP^+^ Luci^+^, EXT1-KO-Rescue GFP^+^ Luci^+^, and EXT1-KO-D164A Rescue GFP^+^ Luci^+^ cells were injected into the mammary fat pad of female NOD-SCID mice (MDA-MB-231(1 × 10^6^) cells. For overexpression MDA-MB-468 WT, GFP^+^ Luci^+^, EXT1 OE GFP^+^ Luci^+^ and EXT1 OE-D164A GFP+ Luci^+^ (5 × 10^6^) cells per mouse). After the injection, the tumor growth kinetics were monitored in real-time using the noninvasive bioluminescence imaging system. We harvested the tumors and weighed them after seven weeks for mice injected with MDA-MB-231 or 12 weeks for those injected with MDA-MB-468 cells. All mouse experiments were carried out under The Hebrew University Institutional Animal Care and Use Committee-approved (IACUC) protocol MD-21-16429-5. In addition, the Hebrew University is certified by the Association for Assessment and Accreditation of Laboratory Animal Care (AAALAC).

### Statistical Analysis

Data are shown as mean ± SD from at least three independent biological experiments. All statistical analyses were performed using the R (version 4.0) or GraphPad Prism (version 8.0) statistical analysis programs. If not indicated in the figure legend, all the p-values were calculated using the unpaired two-tailed Student’s *t*-test. Data distribution was assumed to be normal but this was not formally tested. The significance of the mean comparison is present in each figure.

## Acknowledgments

We thank the members of the Shaul laboratory. This work was supported by the Israel Science Foundation (Grant 1816/16 and 299/21) and the Mizutani Foundation for Glycoscience (grant # 220087). B. Solaimuthu was supported by the Lady Davis Fellowship for postdoctoral researchers at The Hebrew University of Jerusalem. The Genomic Applications Laboratory of the Core Research Facility, The Faculty of Medicine, and The Hebrew University of Jerusalem, Israel, performed the RNA-Seq data analysis. The authors declare no competing financial interests. We would like to thank professor Israel Voldavsky from the Israel Institute of Technology for his support and valuable suggestions.

## Author contribution

Conceptualization and Methodology: B. Solaimuthu and Y.D. Shaul. Investigation: B. Solaimuthu, A. Khatib, A. Karmi, M. Lichtenstein, Formal Analysis: A. Hayashi and M. Tanna. Visualization: B. Solaimuthu and A. Hayashi Writing – original draft: B. Solaimuthu and Y.D. Shaul. Writing – review & editing: B. Solaimuthu and Y.D. Shaul. Funding acquisition: Y.D. Shaul.

---

***Fig. S1: EXT1 expression elevated in aggressive breast cancer cell lines.*** **a** Heparinase III treatment reduces HSPG extracellular levels. MDA-MB-231 cells were treated with increasing heparinase III concentration as indicated for one hour, followed by FACS analysis to determine heparin sulfate levels using the 10E4 antibody. The percentage of cells in each gate is presented. **b** Heparan sulfate biosynthesis pathway. A schematic representation of the HSPG initiation and chain elongation steps in the biosynthesis pathway and their corresponding enzymes. Red arrows indicated the enzymatic reaction outcome. **c** EXT1 expression level is upregulated in mesenchymal cell lines. Cancer cell lines were divided into epithelial (n=378 cell lines) and mesenchymal (n=150 cell lines) groups based on the expression of known mesenchymal markers. Box plots represent the actual expression levels of each gene in the different groups. The p-value was determined by Student’s *t*-test.

***Fig. S2. Validating EXT1 overexpression in mesenchymal cells.*** **a** *EXT1* mRNA level is upregulated in mesenchymal breast cancer cell lines. *EXT1* relative mRNA level was determined for each cell line by qRT-PCR. Each value represents the mean ± SD for n=3. **b** *EXT1* mRNA level is upregulated by the HMLE-Twist-ER-inducible EMT system. Every three days, cells were collected, and mRNA was isolated and subjected to qRT-PCR using the indicated primers. Each value represents the mean ± SD for n=3. **c** *EXT1* expression is significantly correlated with the EMT gene set in different tumor types. patients’ gene expression data from different indicated cancer types that was generated by the TCGA (PanCancer Atlas project) and analyzed using the cBioportal web tool (https://www.cbioportal.org). In these samples, the expression of *EXT1* was compared to the whole transcriptome (~20,000 genes). The genes were then ranked based on the obtained Spearman’s rank correlation coefficient followed and subjected to gene set enrichment analysis (GSEA). GSEA computed the normalized enrichment score (NES) and false discovery rate (FDR) values. **d** EXT1 expression elevates aggressive breast cancer subtypes. Breast cancer patients’ gene expression data from the TCGA (PanCancer Atlas project) database (left) and the METBRIC (right) databases were collected from the cBioportal web tool (https://www.cbioportal.org). The graphs represent *EXT1* expression profiles in different breast cancer subtypes. The p-value was determined by Student’s *t*-test. **e** *EXT1* expression is associated with overall survival (OE). Kaplan-Meier survival plots for patients with breast cancer were divided into high *EXT1* expression (“high” (purple)) and low (“low” (green)). The survival data and the statistics were gained from the METBRIC database and generated using the cBioportal web tool. The p-value was determined by the website.

***Fig. S3. EXT1 overexpression induces cell migration in breast cancer cell lines.*** **a** Both WT and EXT1-FLAG MCF-7 cells were subjected to comparative RNA-Seq analysis. EXT1 demonstrated the highest expression ratio as compared between WT and EXT1-FLAG cells. EXT1 gene expression ratio and the p-value is marked by a blue ball. **b** Representative image of the wound healing scratch assay. Images represent scratch confluency during 0 and 48 h for WT, EXT1-FLAG, and EXT1-D164A-FLAG in MDA-MB-468 cells. Scale bar=400μm. **c** EXT1 overexpression induces the migration capabilities in MCF-7 and MDA-MB-468 cell lines. The migratory capability of the different samples was determined using Boyden chamber-based transwell assay. Representative images are shown after 24 h for WT, EXT1-FLAG, and EXT1-D164A-FLAG for each cell line. The number of migrated cells per 25,000 seeded cells; each value represents the mean ± SD for n=3. Scale bar=100μm. **d** EXT1 overexpression induces the invasion capabilities in MCF-7 and MDA-MB-468 cell lines. The migratory capability of the different samples was determined using Boyden chamber-based transwell assay. Representative images are shown after 24 h for WT, EXT1-FLAG, and EXT1-D164A-FLAG in MCF-7 and MDA-MB-468 cell lines. The number of invaded cells per 20,000 seeded cells. Scale bar=100μm. **e** Heparinase III treatment inhibits the migration ability in EXT1 overexpressed MDA-MB-468 cells. The migratory capability of the different samples was determined using Boyden chamber-based transwell assay. The number of migrated cells per 25,000 seeded cells; each value represents the mean ± SD for n=3. The p-value was determined by Student’s t-test. Scale bar=100μm. **f** EXT1overexpression induces tumor initiation in mice. The tumors developed from EXT1-FLAG cells are larger than those from WT (EV) and EXT1-D164A-FLAG cells. The representative images of tumors were harvested from mice injected with MDA-MB-468 WT (EV) cells (top), EXT1-FLAG cells (middle), and EXT-D164A-FLAG cells (bottom).

***Fig. S4. EXT1 loss inhibits cell migration in breast cancer cell lines.*** **a** Representative image of the scratch assay and confluency during 0 and 22 h for MDA-MB-231 WT, EXT1-KO, EXT1-KO + EXT-FLAG, and EXT1-KO + EXT1-D164A-FLAG cells. Scale bar=400μm. **b** EXT1 loss inhibits MDA-MB-231 cells migration capabilities. The migratory capability of the different samples was determined using Boyden chamber-based transwell assay. Representative Images are shown after 24 h for MDA-MB-231 WT, EXT1-KO, EXT1-KO + EXT-FLAG, and EXT1-KO + EXT1-D164A-FLAG cells. The number of migrated cells per 20,000 seeded cells. Scale bar=100μm. **c** EXT1 loss inhibits the invasion property in the MDA-MB-231 cell line. The migratory capability of the different samples was determined using Boyden chamber-based transwell assay. Representative Images are shown after 24 h for MDA-MB-231 WT, EXT1-KO, EXT1-KO + EXT-FLAG, and EXT1-KO + EXT1-D164A-FLAG cells. The number of invaded cells per 25,000 seeded cells. Scale bar=100μm. **d**. Heparinase III treatment reduces MDA-MB-231 migratory capabilities. MDA-MB-231 cells were treated with heparinase III (10μM). The migratory capability of the different samples was determined using Boyden chamber-based transwell assay. Representative images of each sample (left). Scale bar=100μm. Quantification of data is reported as the number of migrated cells per 20,000 seeded cells (right); each bar represents the mean ± SD for n=3. The p-value was determined by Student’s *t*-test. **e** EXT1 KO in breast cancer cell lines MDA-MB-231 and Hs-578-T cell lines were generated using the CRISPR-Cas9 system. Cells were lysed and subjected to immunoblot using the indicated antibodies. WT= control cells, KO = EXT1 knockout. **f** EXT1 and other EMT marker mRNA expression levels were decreased in the (EXT1-KO) breast cancer cell lines (MDA-MB-231, Hs-575-t). EXT1 expression mRNA level was determined by qRT-PCR. Each value represents the mean ± SD for n=3. **g** EXT1 loss inhibits the migration capabilities in the Hs-578-T cell line. The migratory capability of the different samples was determined using Boyden chamber-based transwell assay. Representative images are shown after 24 h for Hs-578-T WT, and EXT1-KO cells (top). Quantification of data is reported as the number of migrated cells per 20,000 seeded cells (bottom); each bar represents the mean ± SD for n=3. The p-value was determined by Student’s *t*-test. Scale bar=100μm. **h** EXT1 loss inhibits tumor formation in mice. The tumors developed from EXT1-KO cells are smaller than those from EXT1-WT cells. The representative images of tumors were harvested from mice injected with MDA-MB-231 WT cells (top), and EXT-KO cells (bottom).

***Fig. S5: EXT1-KO downregulated the IL-6/STAT3 signaling pathway in breast cancer cells*** **a** *IL-6* expression is significantly reduced in EXT1-KO cells. MDA-MB-231 WT and EXT1-KO cells were subjected to CEL-Seq analysis. The expression ratio of all genes was calculated and ranked based on the relative expression in EXT1 WT and KO. The samples were subjected to GSEA, and the results are presented as a summary plot. **b** EXT1 loss leads to a reduction in the gene expression pattern for “Hallmark of IL6-STAT3 signaling and Hallmark of EMT “. The FDR q-value was computed by GSEA. **c** Heparinase III treatment for 1hr reduced STAT3 phosphorylation in MDA-MB-231 cells. Cells were lysed and subjected to immunoblotting using the indicated antibodies. **d** EXT1 loss induces a global reduction in the expression of IL6 and STAT3 targets. The expression level in WT and EXT1 KO cells is presented in a volcano plot. IL6 (blue) and STAT3 targets (red). **e** CA-STAT3 induces IL-6 expression in MDA-MB-231 cells. RNA was isolated from WT, EXT1-KO, and EXT1-KO + CA-STAT3 cells, and the relative *IL-6* mRNA expression was determined by qRT-PCR. Each value represents the mean ± SD for n = 3. **f** Treating EXT1-FLAG cells with ruxolitinib, inhibited STAT3 phosphorylation in EXT1 overexpressed (EXT1-FLAG -MDA-MB-468) cells were starved with 0% FBS medium for 16 h and treated with or without 10 μM ruxolitinib (JAK1/2 inhibitor) for 12 h. Immunoblots representing MDA-MB-468 cells treated with vehicle control or EXT1-FLAG or 10 μM ruxolitinib for 12 h. Cells were lysed and subjected to immunoblotting using the indicated antibodies.

## Tables

Table S1 shows the RNA-seq analysis of wild-type MCF-7 (VC) and EXT1 overexpressing (EXT1-FLAG).

Table S2 shows the CEL-Seq results of WT and EXT1-KO cells in MDA-MB-231 cells.

## Reference

1. Luo, M., Brooks, M. & Wicha, M. Epithelial-Mesenchymal Plasticity of Breast Cancer Stem Cells: Implications for Metastasis and Therapeutic Resistance. Curr Pharm Design 21, 1301–1310 (2015).

2. Williams, E. D., Gao, D., Redfern, A. & Thompson, E. W. Controversies around epithelial–mesenchymal plasticity in cancer metastasis. Nat Rev Cancer 19, 716–732 (2019).

3. Brabletz, S., Schuhwerk, H., Brabletz, T. & Stemmler, M. P. Dynamic EMT: a multi-tool for tumor progression. Embo J 40, e108647 (2021).

4. Bakir, B., Chiarella, A. M., Pitarresi, J. R. & Rustgi, A. K. EMT, MET, Plasticity, and Tumor Metastasis. Trends Cell Biol 30, 764–776 (2020).

5. Karagiorgou, Z., Fountas, P. N., Manou, D., Knutsen, E. & Theocharis, A. D. Proteoglycans Determine the Dynamic Landscape of EMT and Cancer Cell Stemness. Cancers 14, 5328 (2022).

6. Vallet, S. D., Clerc, O. & Ricard-Blum, S. Glycosaminoglycan–Protein Interactions: The First Draft of the Glycosaminoglycan Interactome. J Histochem Cytochem 69, 93–104 (2020).

7. Hassan, N., Greve, B., Espinoza-Sánchez, N. A. & Götte, M. Cell-surface heparan sulfate proteoglycans as multifunctional integrators of signaling in cancer. Cell Signal 77, 109822 (2020).

8. Onyeisi, J. O. S., Ferreira, B. Z. F., Nader, H. B. & Lopes, C. C. Heparan sulfate proteoglycans as targets for cancer therapy: a review. Cancer Biol Ther 21, 1087–1094 (2020).

9. Marques, C., Reis, C. A., Vivès, R. R. & Magalhães, A. Heparan Sulfate Biosynthesis and Sulfation Profiles as Modulators of Cancer Signalling and Progression. Frontiers Oncol 11, 778752 (2021).

10. Esko, J. D. & Selleck, S. B. ORDER OUT OF CHAOS: Assembly of Ligand Binding Sites in Heparan Sulfate1. Annu Rev Biochem 71, 435–471 (2002).

11. Couchman, J. R. Syndecan-1 (CD138), Carcinomas and EMT. Int J Mol Sci 22, 4227 (2021).

12. Ibrahim, S. A. et al. Syndecan-1 is a novel molecular marker for triple negative inflammatory breast cancer and modulates the cancer stem cell phenotype via the IL-6/STAT3, Notch and EGFR signaling pathways. Mol Cancer 16, 57 (2017).

13. Rmaileh, A. A. et al. DPYSL2 interacts with JAK1 to mediate breast cancer cell migration. J Cell Biol 221, e202106078 (2022).

14. Sarrazin, S., Lamanna, W. C. & Esko, J. D. Heparan Sulfate Proteoglycans. Csh Perspect Biol 3, a004952 (2011).

15. Poulain, F. E. & Yost, H. J. Heparan sulfate proteoglycans: a sugar code for vertebrate development? Development 142, 3456–3467 (2015).

16. Li, J.-P. & Kusche-Gullberg, M. Chapter Six Heparan Sulfate: Biosynthesis, Structure, and Function. Int Rev Cel Mol Bio 325, 215–273 (2016).

17. Busse-Wicher, M., Wicher, K. B. & Kusche-Gullberg, M. The extostosin family: Proteins with many functions. Matrix Biol 35, 25–33 (2014).

18. McCormick, C., Duncan, G., Goutsos, K. T. & Tufaro, F. The putative tumor suppressors EXT1 and EXT2 form a stable complex that accumulates in the Golgi apparatus and catalyzes the synthesis of heparan sulfate. Proc National Acad Sci 97, 668–673 (2000).

19. Busse, M. et al. Contribution of EXT1, EXT2, and EXTL3 to Heparan Sulfate Chain Elongation*. J Biol Chem 282, 32802–32810 (2007).

20. Cheung, P. K. et al. Etiological Point Mutations in the Hereditary Multiple Exostoses Gene EXT1: A Functional Analysis of Heparan Sulfate Polymerase Activity. Am J Hum Genetics 69, 55–66 (2001).

21. Stickens, D., Zak, B. M., Rougier, N., Esko, J. D. & Werb, Z. Mice deficient in Ext2 lack heparan sulfate and develop exostoses. Development 132, 5055–5068 (2005).

22. Bukowska-Olech, E. et al. Hereditary Multiple Exostoses—A Review of the Molecular Background, Diagnostics, and Potential Therapeutic Strategies. Frontiers Genetics 12, 759129 (2021).

23. David, G., Bai, X. M., Schueren, B. V. der, Cassiman, J. J. & Berghe, H. V. den. Developmental changes in heparan sulfate expression: in situ detection with mAbs. J Cell Biology 119, 961–975 (1992).

24. Yang, J. et al. Twist, a Master Regulator of Morphogenesis, Plays an Essential Role in Tumor Metastasis. Cell 117, 927–939 (2004).

25. Tam, W. L. et al. Protein Kinase C α Is a Central Signaling Node and Therapeutic Target for Breast Cancer Stem Cells. Cancer Cell 24, 347–364 (2013).

26. Ren, Z. et al. Syndecan-1 promotes Wnt/β-catenin signaling in multiple myeloma by presenting Wnts and R-spondins. Blood 131, 982–994 (2018).

27. Shaul, Y. D. et al. MERAV: a tool for comparing gene expression across human tissues and cell types. Nucleic Acids Res 44, D560–D566 (2016).

28. Shaul, Y. D. et al. Dihydropyrimidine Accumulation Is Required for the Epithelial-Mesenchymal Transition. Cell 158, 1094–1109 (2014).

29. Kanehisa, M., Goto, S., Kawashima, S. & Nakaya, A. The KEGG databases at GenomeNet. Nucleic acids research 30, 42–46 (2002).

30. Kanehisa, M., Goto, S., Sato, Y., Furumichi, M. & Tanabe, M. KEGG for integration and interpretation of large-scale molecular data sets. Nucleic acids research 40, D109–14 (2012).

31. Mani, S. A. et al. The Epithelial-Mesenchymal Transition Generates Cells with Properties of Stem Cells. Cell 133, 704–715 (2008).

32. Gao, J. et al. Integrative Analysis of Complex Cancer Genomics and Clinical Profiles Using the cBioPortal. Sci Signal 6, pl1–pl1 (2013).

33. Ciriello, G. et al. Comprehensive Molecular Portraits of Invasive Lobular Breast Cancer. Cell 163, 506–519 (2015).

34. Curtis, C. et al. The genomic and transcriptomic architecture of 2,000 breast tumours reveals novel subgroups. Nature 486, 346–352 (2012).

35. Subramanian, A. et al. Gene set enrichment analysis: A knowledge-based approach for interpreting genome-wide expression profiles. P Natl Acad Sci Usa 102, 15545–15550 (2005).

36. Györffy, B. et al. An online survival analysis tool to rapidly assess the effect of 22,277 genes on breast cancer prognosis using microarray data of 1,809 patients. Breast cancer research and treatment 123, 725–731 (2010).

37. Solaimuthu, B., Hayashi, A., Khatib, A. & Shaul, Y. D. Monitoring Breast Cancer Growth and Metastatic Colony Formation in Mice using Bioluminescence. J Vis Exp (2021) doi:10.3791/63060.

38. Hashimshony, T. et al. CEL-Seq2: sensitive highly-multiplexed single-cell RNA-Seq. Genome Biol 17, 77 (2016).

39. Fischer, M. et al. A bioactive designer cytokine for human hematopoietic progenitor cell expansion. Nat Biotechnol 15, 142–145 (1997).

40. Wang, Y., Shen, Y., Wang, S., Shen, Q. & Zhou, X. The role of STAT3 in leading the crosstalk between human cancers and the immune system. Cancer Lett 415, 117–128 (2018).

41. Khatib, A. et al. The glutathione peroxidase 8 (GPX8)/IL-6/STAT3 axis is essential in maintaining an aggressive breast cancer phenotype. Proc National Acad Sci 117, 21420–21431 (2020).

42. Jiang, G.-X. et al. Interleukin-6 induces epithelial-mesenchymal transition in human intrahepatic biliary epithelial cells. Mol Med Rep 13, 1563–1569 (2016).

43. Gyamfi, J., Lee, Y.-H., Eom, M. & Choi, J. Interleukin-6/STAT3 signalling regulates adipocyte induced epithelial-mesenchymal transition in breast cancer cells. Scientific Reports 8, 1–13 (2018).

44. Martz, C. A. et al. Systematic identification of signaling pathways with potential to confer anticancer drug resistance. Science signaling 7, ra121 (2014).

45. Mesa, R. A. Ruxolitinib, a selective JAK1 and JAK2 inhibitor for the treatment of myeloproliferative neoplasms and psoriasis. Idrugs Investigational Drugs J 13, 394–403 (2010).

46. Ibrahim, S. A. et al. Syndecan-1 (CD138) Modulates Triple-Negative Breast Cancer Stem Cell Properties via Regulation of LRP-6 and IL-6-Mediated STAT3 Signaling. Plos One 8, e85737 (2013).

47. Sayyad, M. R. et al. Syndecan-1 facilitates breast cancer metastasis to the brain. Breast Cancer Res Tr 178, 35–49 (2019).

48. Qiao, W., Liu, H., Guo, W., Li, P. & Deng, M. Prognostic and clinical significance of syndecan-1 expression in breast cancer: A systematic review and meta-analysis. Eur J Surg Oncol 45, 1132–1137 (2019).

49. Porter, D. E. et al. Severity of disease and risk of malignant change in hereditary multiple exostoses. A genotype-phenotype study. J Bone Jt Surg Br Volume 86, 1041–6 (2004).

50. Alvarez, C. M., Vera, M. A. D., Heslip, T. R. & Casey, B. Evaluation of the Anatomic Burden of Patients with Hereditary Multiple Exostoses. Clin Orthop Relat R 462, 73–79 (2007).

51. Daakour, S. et al. Systematic interactome mapping of acute lymphoblastic leukemia cancer gene products reveals EXT-1 tumor suppressor as a Notch1 and FBWX7 common interactor. Bmc Cancer 16, 335 (2016).

52. Dong, S. et al. Increased EXT1 gene copy number correlates with increased mRNA level predicts short disease-free survival in hepatocellular carcinoma without vascular invasion. Medicine 97, e12625 (2018).

53. Julien, S. et al. Selectin Ligand Sialyl-Lewis x Antigen Drives Metastasis of Hormone-Dependent Breast Cancers. Cancer Res 71, 7683–7693 (2011).

54. Taghavi, A. et al. Gene expression profiling of the 8q22-24 position in human breast cancer: TSPYL5, MTDH, ATAD2 and CCNE2 genes are implicated in oncogenesis, while WISP1 and EXT1 genes may predict a risk of metastasis. Oncol Lett 12, 3845–3855 (2016).

55. Manandhar, S. et al. Exostosin 1 regulates cancer cell stemness in doxorubicin-resistant breast cancer cells. Oncotarget 5, 70521–70537 (2017).

56. Kreuger, J. & Kjellén, L. Heparan Sulfate Biosynthesis. J Histochem Cytochem 60, 898–907 (2012).

57. Xie, M. & Li, J. Heparan sulfate proteoglycan – A common receptor for diverse cytokines. Cell Signal 54, 115–121 (2019).

58. Yamada, S. et al. Embryonic Fibroblasts with a Gene Trap Mutation in Ext1 Produce Short Heparan Sulfate Chains*. J Biol Chem 279, 32134–32141 (2004).

59. Wu, T. et al. clusterProfiler 4.0: A universal enrichment tool for interpreting omics data. Innovation 2, 100141 (2021).

60. Győrffy, B. Survival analysis across the entire transcriptome identifies biomarkers with the highest prognostic power in breast cancer. Comput Struct Biotechnology J 19, 4101–4109 (2021).

