## Supplementary figures and images for "The metabolic enzyme EXT1 is sufficient to induce the epithelial-mesenchymal transition program in cancers"

### Supplemental figures

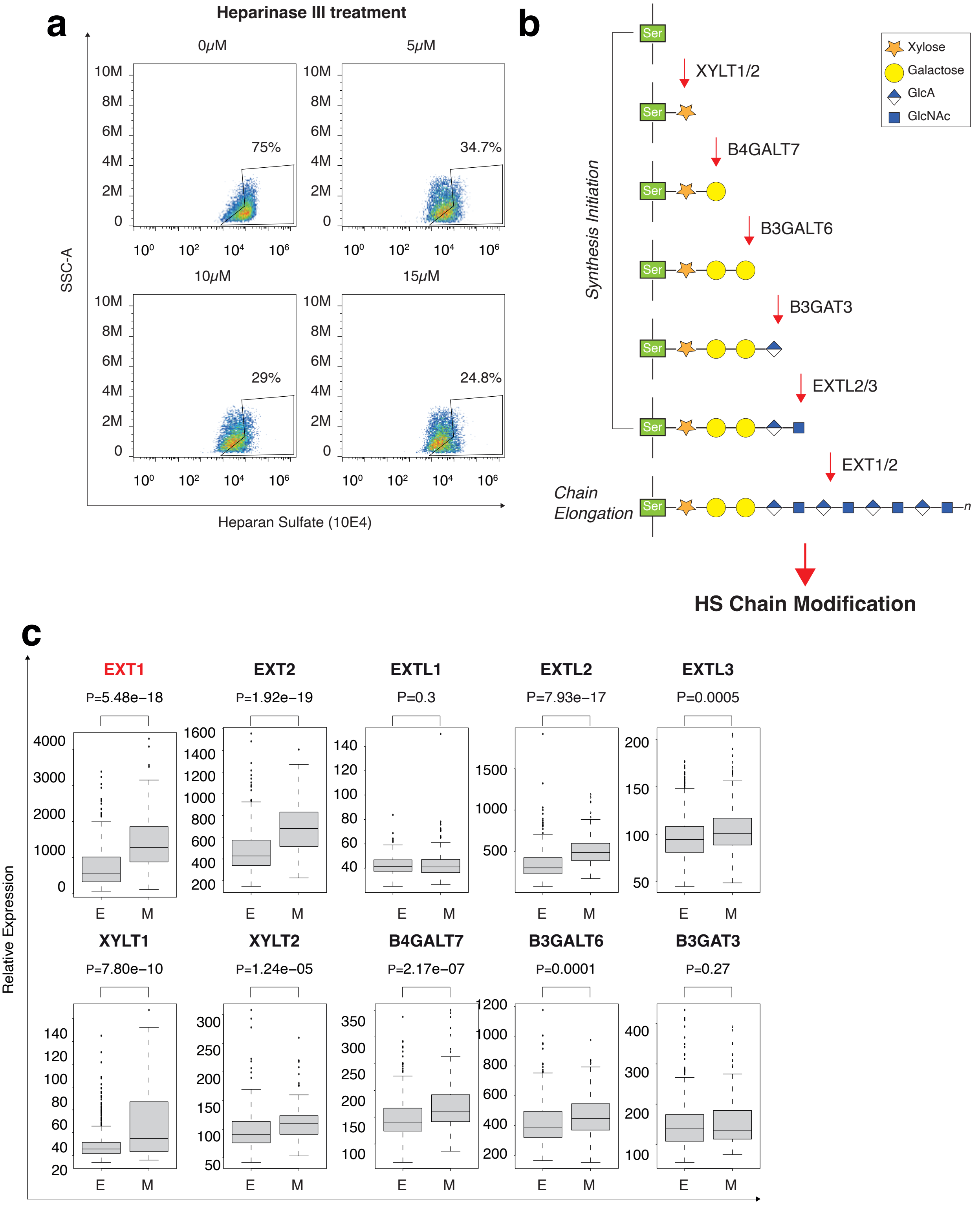

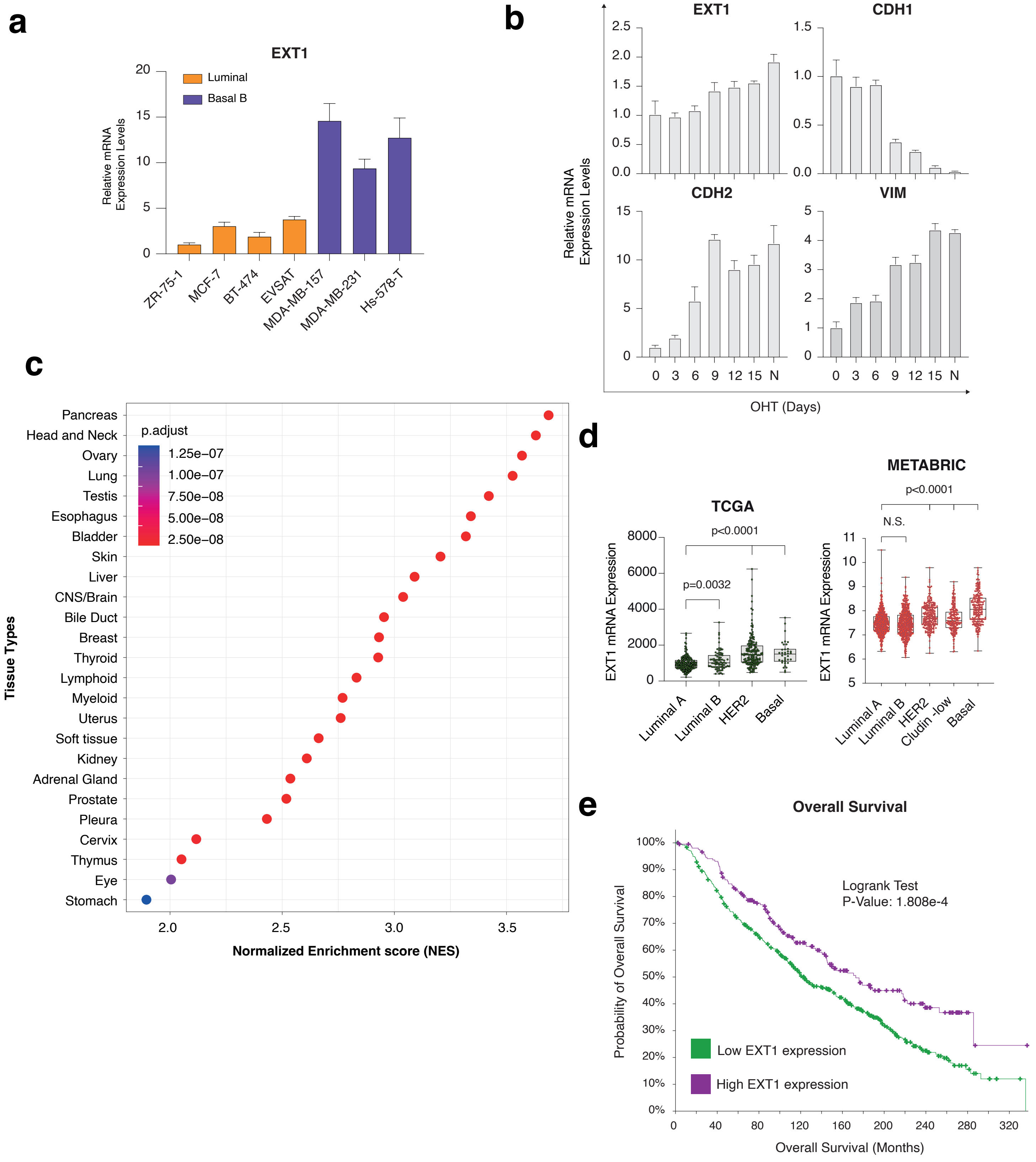

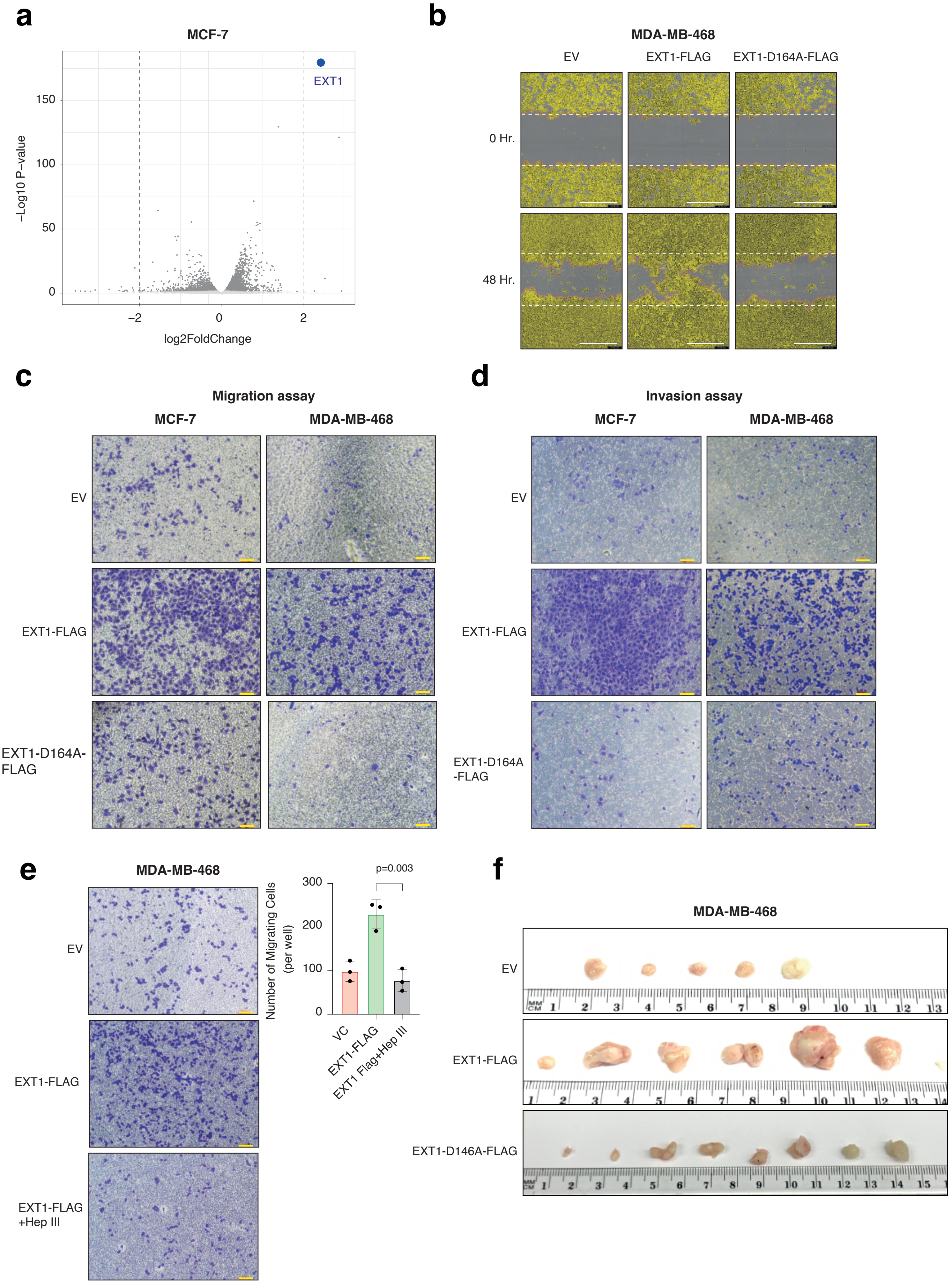

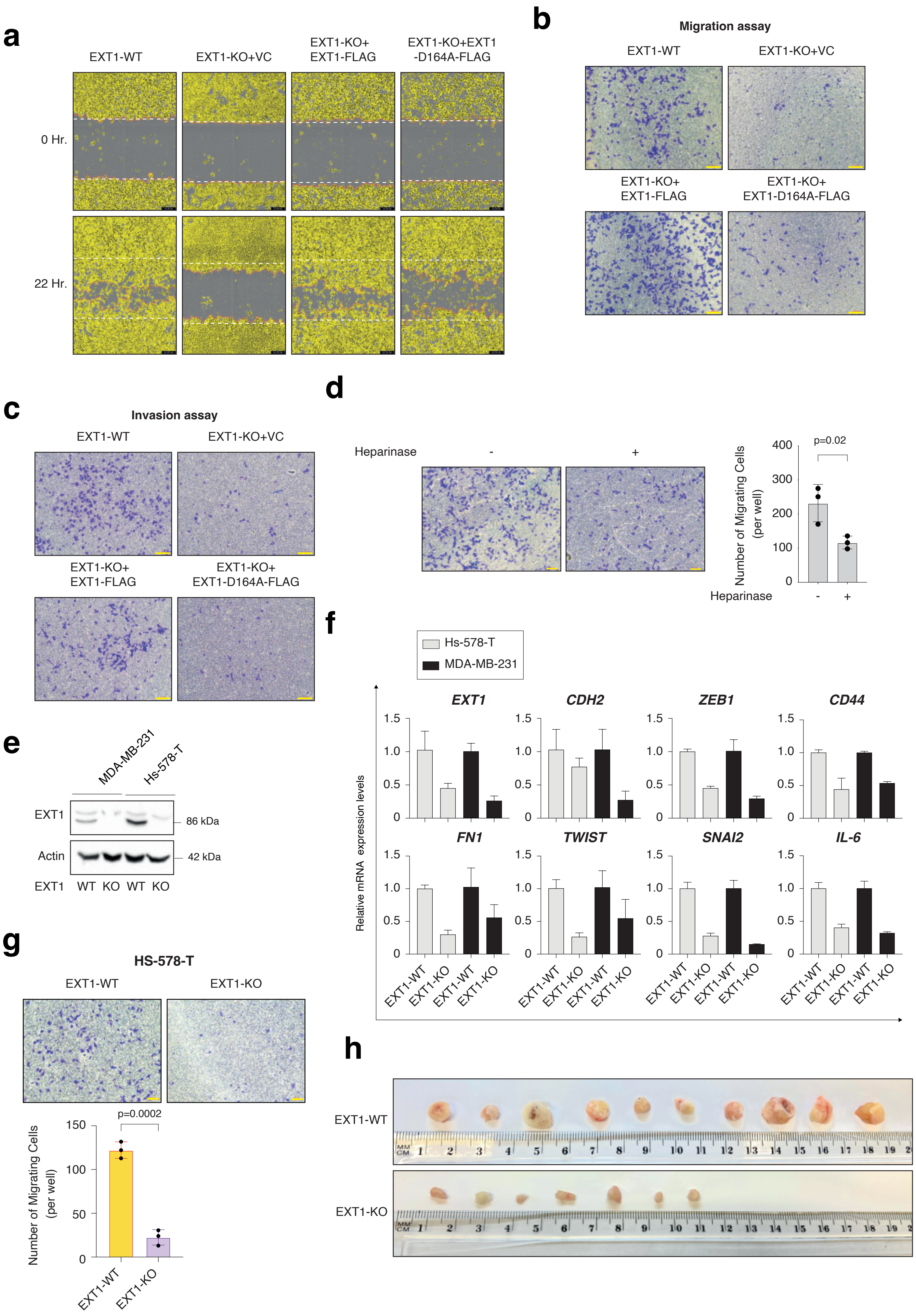

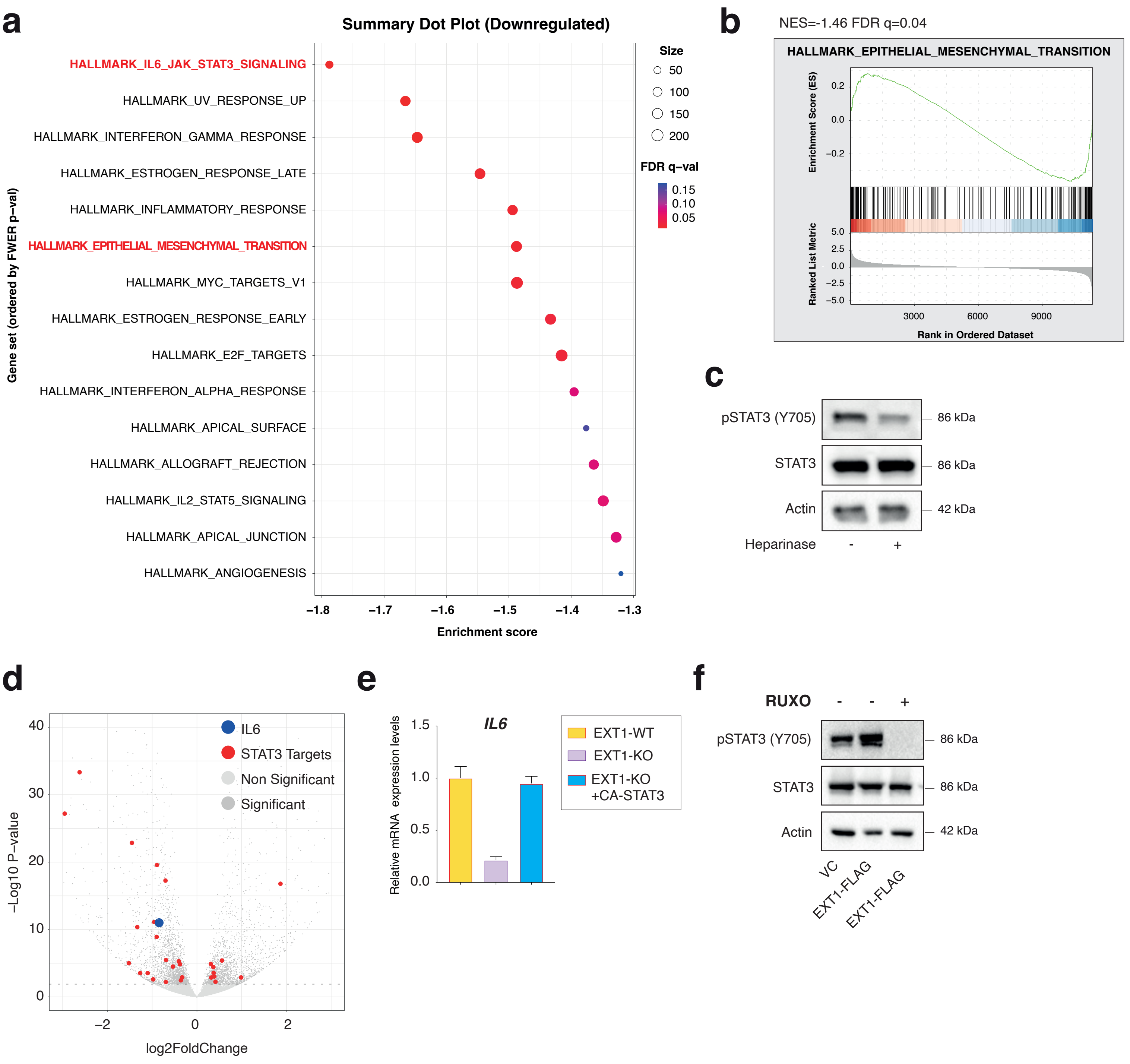
